# The *Verticillium longisporum* phospholipase VlsPLA_2_ is a virulence factor targets host nuclei and modulates plant immunity

**DOI:** 10.1101/2022.03.19.484916

**Authors:** Vahideh Rafiei, Heriberto Vélëz, Edoardo Piombo, Mukesh Dubey, Georgios Tzelepis

## Abstract

Phospholipases A_2_ (PLA_2_) are lipolytic enzymes, responsible for phospholipids hydrolysis. The role of PLA_2_ in various aspects of cell physiology has been shown, but their involvement in host-microbe interactions remains to be elucidated. The present study investigated the biological function of the secreted VIsPLA_2_ phospholipase in the phytopathogenic fungus *Verticillium longisporum* with emphasis on its role in host-microbe interactions. The *VlsPLA*_*2*_ was highly induced in *V. longisporum* during interaction with host plant *Brassica napus*, encoding an active phospholipase A_2_. VlsPLA_2_-overexpression *V. longisporum* strain showed an increased virulence on *Arabidopsis* plants, plausibly by inducing virulence factors. Furthermore, VIsPLA_2_ are transported to the nucleus, by hijacking VAMPA proteins, causing suppression of PTI-induced hypersensitive response, possibly by modulating the expression of genes involved in plant immunity. In summary, VlsPLA_2_ acts as a virulence factor by hydrolyzing the hosts nuclear envelope phospholipids, an action that induces signaling cascade, suppressing basal plant immunity responses.

## Introduction

Phospholipids are essential structural components of plasma membranes, being conserved in living organisms, from bacteria to plants and humans. They form a lipid bilayer with a hydrophobic and hydrophilic environment and are abundant in mitochondrial and nuclear envelopes, whereas they are scarce in the thylakoid membranes^1^. Phospholipases are enzymes that hydrolyze phospholipids into phosphatidic acid (PA), diacylglycerol (DAG), free fatty acids (FFAs) and lysophospholipids (LPLs). Based on the site of glycerophospholipid hydrolysis, phospholipases are categorized in phospholipases A_1_ (PLA_1_) and A_2_ (PLA_2_); phospholipases C (PLC); phospholipases D (PLD), and are further subdivided in many families and subfamilies^2^. They play important roles in different aspects of cell physiology including signal transduction, cytoskeletal dynamics, and protein secretion^3^. They are also involved in plant defense mechanisms probably as early signaling molecules^4^. Previous studies showed that secreted phospholipases are involved in the regulation of different processes in bacterial and fungal pathogens, but their precise roles in fungal virulence remain unknown^5,6^.

Pathogens secrete a plethora of small proteins, termed effectors, in order to avoid plant immunity mechanisms and to establish a successful infection^7^. These proteins can have diverse functions: for example to induce necrosis; to suppress the hypersensitive response (HR); to protect fungal hyphae from host secreting enzymes, such as chitinases; or can be involved in host energy production^8^. Although the sequencing of fungal genomes has helped to identify and understand the diverse roles of fungal effectors, there are still many aspects that remain to be elucidated. From the plant side, the host deploys defense mechanisms to counteract the pathogen’s attack and hence, signaling plays a prominent role in this process. As the first layer, the plasma membrane-localized pattern recognition receptors (PRRs) recognize the microbe associated molecular patterns (MAMPs) (e.g., chitin, flagellin, etc.) and this recognition can lead to a burst of reactive oxidate species (ROS), influx of calcium (Ca^+^), the activation of mitogen-activated protein kinases (MAPK) and the induction of defense genes, resulting in pattern triggered immunity (PTI)^9^. When pathogens overcome PTI, then hosts deploy Nod-like receptors (NLRs) able to recognize effectors, leading to effector triggered immunity (ETI)^10^. The outcome of ETI is usually development of HR, although PTI can also result in program cell death^11^.

The genus *Verticillium*, which belongs to division Ascomycota in the Plectosphaerellaceae family, includes species causing mainly vascular wilt disease in a plethora of economically important crops^12^. The most important one is the haploid species, *V. dahliae*, which has a broad host range comprising a high number of dicotyledonous plants (e.g., tomato, cotton, olive trees, etc.) ^12^. In contrast, the *V. longisporum* species has a narrow host range, infecting mostly plants in Brassicaceae family (e.g., rape seed, cabbage, and broccoli)^13^, and is one of the most prominent pathogens in rape seed (*Brassica napus*) cultivation worldwide^13,14,15^. *Verticillium longisporum* is the only non-haploid species in this genus and its genome, characterized as amphidiploid, is the result of hybridization of two species, *V. dahliae* and an unknown lineage^16,17,18^. Progenitors were named A1, D1, D2, and D3, leading to three *V. longisporum* lineage compositions A1/D1, A1/D2 and A1/D3. The origin of A1 and D1 progenitors is unknown, while D2 and D3 derived from two *V. dahliae* lineages^17,18^

In this study, we investigated the role of the secreted phospholipase VlsPLA_2_ in virulence of *V. longisporum. VlsPLA*_*2*_ was highly induced during the interaction with *B. napus*. Overexpression of this gene in *V. longisporum* resulted in mutants with increased fungal virulence, and upregulation of genes coding for variable pathogenicity factors. Furthermore, we confirmed that this protein is an active PLA_2_, that seemed to interact with vesicle associated membrane proteins (VAMPs), which are essential compartments of plasma membranes and involved in cellular trafficking ^19,20^. This interaction facilitates VlsPLA_2_ facilitating its transfer to the nuclear envelope and entry to the nucleoplasm, where it altered the expression of genes involved in plant defenses. Lastly, VlsPLA_2_ was able to suppress the PTI-related HR, induced by the Cf4/Avr4 complex. Taken together, the results from the current study provide a better picture towards understanding the precise roles that these lytic enzymes play in fungal infection biology.

## Results

### The *VlsPLA*_*2*_ gene is highly induced during host infection

*Verticillium longisporum* VL1 isolate contains more than 80 genes encoding putative candidate effector proteins^21^. Since the up-regulation of genes during host infection is a criterion used to classify the encoded protein as an effector^8^, we prioritized 13 predicted singletons from the VL1 genome and studied their transcription patterns during infection. Among these genes, only seven showed to be expressed during interaction with the host. The *VL1_T00014035* gene, encoding a putative phospholipase A_2_ (denominated as VlsPLA_2_), was highly upregulated in *V. longisporum* during interaction with *B. napus* roots compared to the mycelium, grown in potato dextrose broth (PDB). (Supplementary Fig. S1). The *VlsPLA*_2_ gene transcription pattern showed the highest induction at six days post infection (Supplementary Fig. S1). Since the role of secreted A_2_ phospholipases in fungal virulence are understudied, this gene was selected for further analysis.

### VlsPLA_2_ is a functional phospholipase A_2_ enzyme

Analysis of *V. longisporum* genome identified two alleles of the *VlsPLA*_*2*_ gene (*VL1_T00014035* and *VL1_T00011351)*. Conserved domain analysis using translated amino acid sequences of VlsPLA_2_ identified a conserved phospholipase A_2_ domain (IPR015141), known to be present in fungal and bacterial phospholipase A_2_ enzymes^22,23^. Furthermore, sequence analysis of VlsPLA_2,_ derived from fungal and bacterial species, showed a conserved CxRHDFGYxN motif, including the catalytic dyad (HD) (Fig. 1a). Further analysis with multiple prediction tools, showed that VlsPLA_2_ was a putative cytoplasmic effector, while the presence of a chloroplast transit peptide (cTP) and two nuclear localization signals (NLS; one monopartite and one bipartite), were predicted, indicating a nuclear and chloroplast localization of this effector (Fig. 1a).

**Fig. 1.**
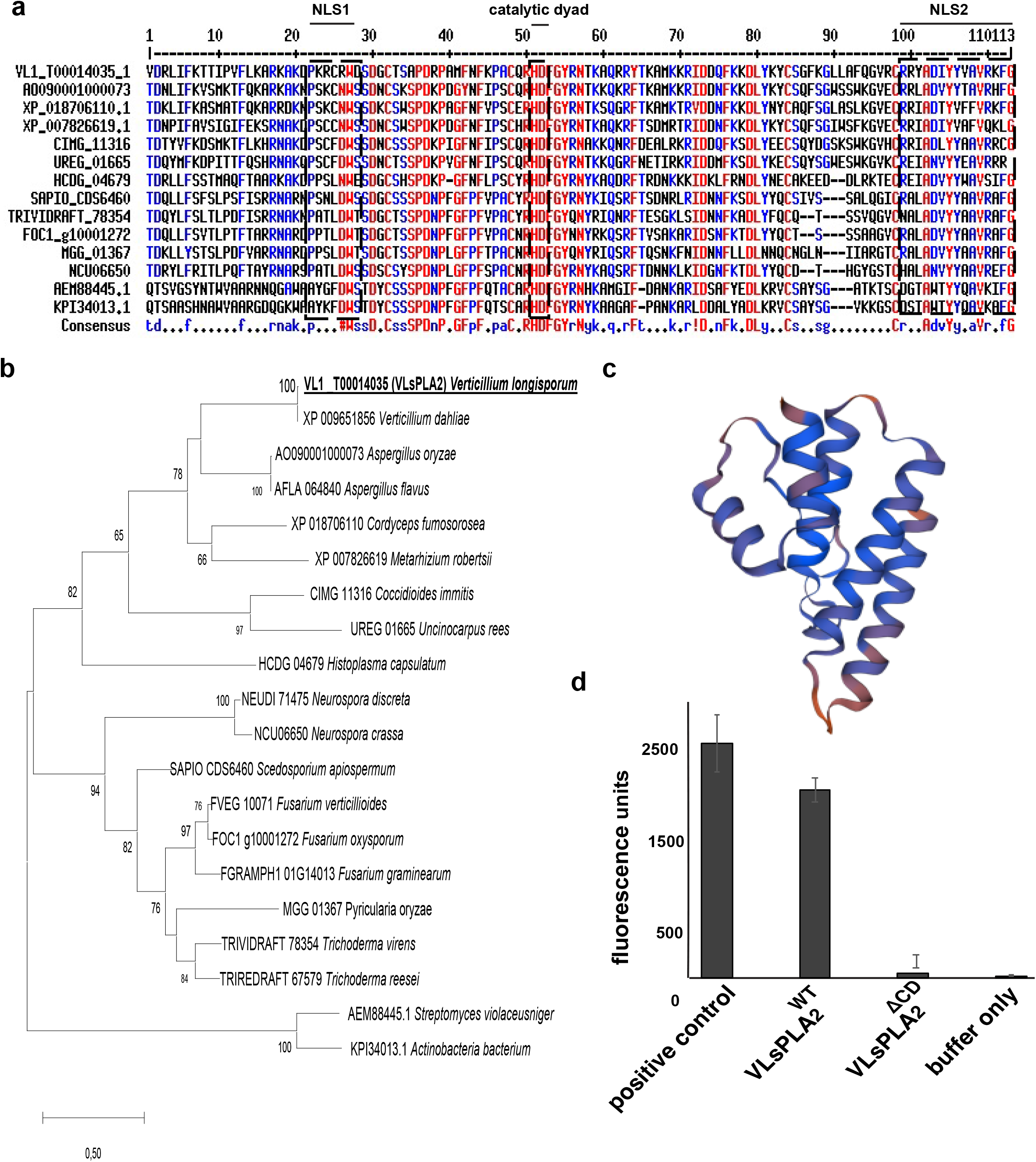
Analysis of the VlsPLA_2_ protein. **a**. Alignment of the amino acid sequences of VlsPLA_2_ and phospholipases derived from different bacterial and fungal species using the Clustal Omega software. Identical amino acids are marked with red color. Different motifs showed in red dashed boxes. The catalytic dyad HD and the two NLS are marked with dashed boxes. **b**. Phylogenetic analysis of the VlsPLA_2_ protein and phospholipases from different fungal species. Analysis was conducted using the neighbour-joining method with the JTT amino acid substitution model based on amino acid sequences and 1000 bootstraps. Number at nodes indicate the bootstrap values. Bar indicates the number of amino acid substitution. Predicted amino acidic sequences were aligned using the CLUSTAL W algorithm and phylogeny was constructed in the MEGA X software. **c**. Predicted 3D structure of VlsPLA_2_. The protein model was predicted using the SWISS-MODEL software. **d**. Phospholipase A_2_ activity assay using 10μM purified VlsPLA_2_^WT^ and VlsPLA2^ΔCD^ proteins. PLA_2_ from bee venom and buffer-only were used as a positive and negative controls respectively. Error bars represent SE based on eight technical replicates.

To study the gene distribution of *PLA*_*2*_ among fungal species, more than 60 genomes from species in Basidiomycota, Ascomycota, Mucoromycota and Chytridiomycota divisions were screened. Our analysis showed that genes, putatively encoding secreted PLA_2_, were identified only in Ascomycota species with diverse lifestyles, and especially in classes Sordariomycetes, Dothideomycetes, Eurotiomycetes and Pezizomycetes, (Supplementary Table S1). A phylogenetic analysis on putative PLA_2_ homologs derived from different fungal species, showed that these proteins grouped in two major clades, and that VlsPLA_2_ was clustered together with PLA_2_ from *V. dahliae, Aspergillus* spp. and entomopathogenic species, such as *Metarhizium* and *Cordyceps* (Fig. 1b). Analysis of the predicted 3D protein structure revealed 54% identity with the PLA_2_ from the mycorrhizal ascomycete *Tuber borchii* and 48% similarity with the PLA_2_ from the bacterial species *Streptomyces violaceoruber*^22,23^ (Fig. 1c).

To investigate the phospholipase activity, VlsPLA_2_ was heterologously expressed in *E. coli* cells, tagged with 6xHis epitope, and purified through a His-tag purification resin. Enzymatic assays with the purified protein showed that VlsPLA_2_ was an active phospholipase A_2_ that can degrade the PLA_2_ selective substrate, similar to the PLA_2_ derived from the venom of *Apis mellifera* that was used as a positive control (Fig. 1d). The role of HD catalytic dyad in enzymatic activity of VlsPLA_2_^24^ was investigated by replacing both amino acids with Alanine. Our results showed that the mutated version of VlsPLA_2_ (i.e., VlsPLA_2_^ΔCD^) was not able to degrade the selective substrate, confirming the importance of this dyad in its enzymatic function (Fig. 1d).

### Heterologous expression of VlsPLA_2_ increases the amount of certain phospholipids in plants

Since phospholipases are the major factor for phospholipid production in cells, we conducted a lipidomic analysis in order to investigate whether VlsPLA_2_ was able to change the phospholipid profile in the plant. Thus, the WT and mutated version of this protein (i.e., VlsPLA2^WT^ and VlsPLA_2_^ΔCD^, respectively), were heterologously expressed in *Nicotiana benthamiana* plants and the lipid profiles were analysed using high resolution Quadruple-Time-Of-flight (QTOF) mass spectrometry. We focused our analysis mostly on classes of phospholipids (i.e., phosphatidylcholine (PC) and phosphatidylethanolamine (PE)), which are abundant in plant plasma membranes^1^. Our data showed that the relative amounts of monoacyl-glycero-phosphocholines (LPC) 16:0, and diacyl-glycero-phosphocholines (PC) 36:5 and 36:6, were significantly increased in plants expressing the VlsPLA_2_^WT^, as compared to the VlsPLA_2_^ΔCD^ or mock-inoculated ones (Supplementary Fig. S2), indicating that this phospholipase has an impact in the hosts phospholipid profile.

### The *VlsPLA*_*2*_ phospholipase contributes to the *V. longisporum* virulence

To evaluate the potential involvement of VlsPLA_2_ in *V. longisporum* virulence, the construction of a deletion strain was attempted. Interestingly, all efforts to delete this gene, either in *V. longisporum* or in *V. dahliae*, resulted in mutants carrying both, the deleted locus, and the WT gene (data not shown). Attempts to segregate the mutated nucleus via multiple single spore isolations, continuously failed. Thus instead, mutants overexpressing the *VlsPLA*_*2*_ gene (constitutively expressed with the *gpdA* promoter from *Aspergillus nidulans*), in *V. longisporum* were generated. The transcription analysis of six single spore isolates, three overexpressing the WT (i.e., *VlsPLA*_*2*_*+*^*WT*^) and three the enzymatically inactive gene (i.e., *VlsPLA*_*2*_*+*^*ΔCD*^), showed a significantly increase of *VlsPLA*_*2*_ transcription levels in comparison to WT (Supplementary Fig. 3). The Isolates *VlsPLA*_*2*_*+*^*WT*^ (3) and *VlsPLA*_*2*_*+*^*ΔCD*^ (1) were chosen for further analysis, since they showed similar expression levels (Supplementary Fig. 3).

*Arabidopsis thaliana* plants were inoculated with the *V. longisporum* WT or overexpression strains, and plant growth and disease symptoms were monitored 28 dpi. We observed that plants infected with the *VlsPLA*_*2*_*+*^*WT*^ strain showed more severe symptoms and smaller rosette, as compared to plants infected with the WT and *VlsPLA*_*2*_*+*^*ΔCD*^ strains. A higher number of dead plants were observed upon infection with the *VlsPLA*_*2*_*+*^*WT*^ strain, as compared to the *VlsPLA*_*2*_*+*^*ΔCD*^ one (Fig. 2a, b, c).

**Fig. 2.**
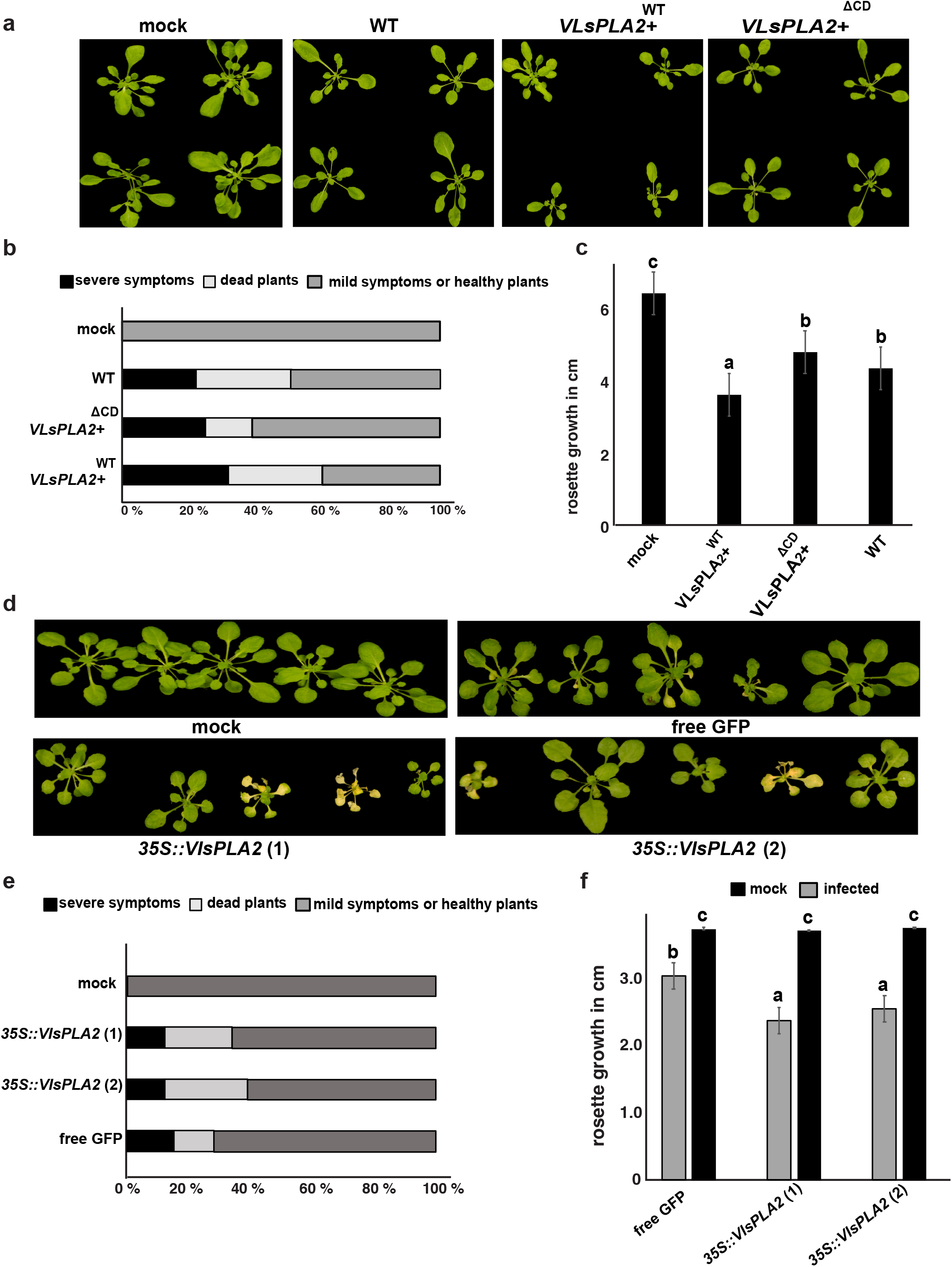
The VlsPLA_2_ phospholipase contributes to *V. longisporum* virulence. **a**. *A. thaliana* plants infected with *V. longisporum* VlsPLA_2_+^WT^, VlsPLA_2_+^ΔCD^, or the wild type (WT) strains. The experiment was performed in 36 plants. Photographs of representative plants were taken 28 days post inoculation (dpi). Mock-inoculated plants were used as control. **b**. percentage of dead, mild-infected and healthy *A. thaliana* plants 28 dpi with *V. longisporum* VlsPLA_2_ +^WT^, VlsPLA +^ΔCD^ or WT strains. **c**. Rosette growth in cm of *A. thaliana* plants infected with *V. longisporum* VlsPLA_2_+^WT^, VlsPLA_2_+^ΔCD^, or WT strains 28 dpi. Different letters (a, b, c) showed statistically significant differences according to Student’s T test (p<0.05). Error bars represents SE based on at least 36 plants. **d**. Representative *A. thaliana* plants, expressing the VlsPLA_2_ phospholipase (*35S::VlsPLA*_*2*_), 28 days post infection with *V. longisporum*. Mock-inoculated and plants expressing free GFP were used as controls. **e**. Percentage of dead, mild-infected and healthy *35S::VlsPLA*_*2*_, free GFP-expressed and mock-inoculated *A. thaliana* plants, 28 dpi with *V. longisporum*. **f**. Rosette growth in cm, of *35S::VlsPLA*_*2*_, free GFP-expressed and mock-inoculated *A. thaliana* plants, 28 dpi with *V. longisporum*. Different letters (a, b, c) showed statistically significant differences according to Student’s T test (p<0.05). Error bars represents SE based on at least 50 plants. Images background has been removed using the Affinity Designer software, without any modification in plants.

Furthermore, the role of this phospholipase in virulence was investigated by generating *A. thaliana* lines overexpressing the *VlsPLA*_*2*_ gene. Our results showed that the overexpression lines (*35S::VlsPLA*_*2*_) displayed higher percentage of dead plants and smaller rosette, compared to the free GFP infected plants, indicating higher susceptibility to *V. longisporum* infection compared to the plants that expressed only free GFP (Fig. 2d, e, f). These data indicate that VlsPLA_2_ has an important contribution to the *V. longisporum* virulence.

Since the *VlsPLA*_*2*_ was highly expressed in *V. longisporum* during interaction with *B. napus* (Supplementary Fig. 1), we analyzed the transcriptome of the *VlsPLA*_*2*_*+*^*WT*^ strain to investigate the potential transcriptomic changes that the upregulation of this gene could cause in *V. longisporum* mycelia during infection. Our data showed that more than 1300 genes were differentially regulated in the *VlsPLA*_*2*_*+*^*WT*^ strain, as compared to the WT (Fig. 3a; Supplementary Table S2). To test whether these transcriptional changes in *VlsPLA*_*2*_*+*^*WT*^ strain was associated with its enhanced phospholipase activity, transcriptome of *VlsPLA*_*2*_*+*^*ΔCD*^ strain was also analyzed. In contrast to the *VlsPLA*_*2*_*+*^*WT*^ strain, only 104 genes were differentially expressed in the *VlsPLA*_*2*_*+*^*ΔCD*^ strain compared to the WT (Fig. 3a; Supplementary Table S2). Moreover, among these 1300 genes, 158 were up regulated in the *VlsPLA*_*2*_^*+WT*^ strain as compared to both the *VlsPLA*_*2*_*+*^*ΔCD*^ and WT (Fig. 3a). Among them, we identified several genes that had been validated for their role in *Verticillium* virulence^25,26,27^, such as genes encoding zinc finger proteins, cytochrome monooxygenases, efflux-pump proteins, and a Calpain-A like protein (Fig. 3b; Supplementary table S3), indicating that *VlsPLA*_*2*_ could be a co-regulator of pathogenicity factors in *V. longisporum*.

**Fig. 3.**
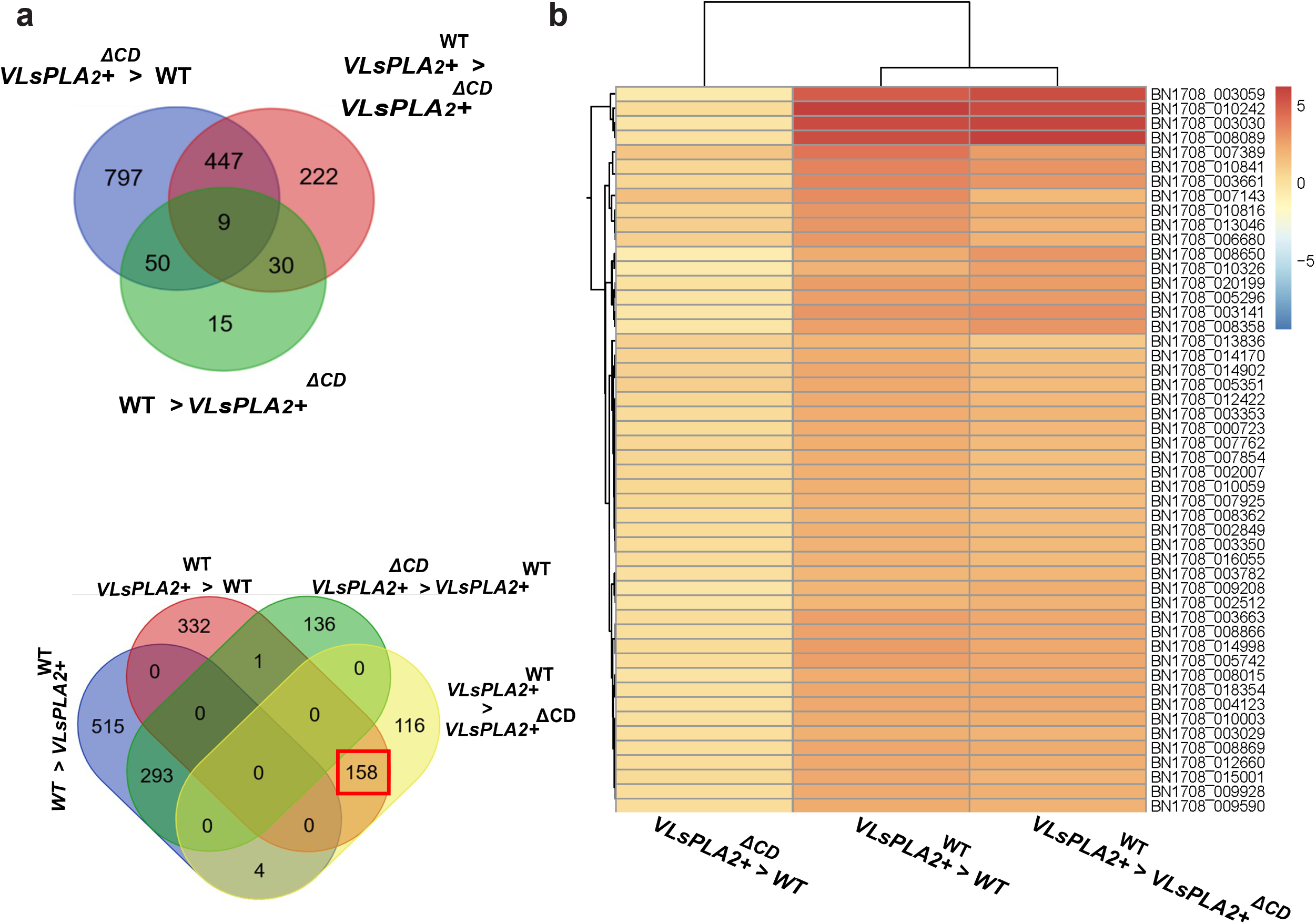
Overexpression of *VlsPLA*_*2*_ causes severe transcriptomic changes in *V. longisporum* mycelia. **a**. Venn diagrams show the number of differentially expressed genes in *V. longisporum* mycelia derived from the WT, and the strains overexpressing either the functionally active (VlsPLA +^WT^) or the inactive (VlsPLA +^ΔCD^) VlsPLA_2_ phospholipase. The sign ‘‘>‘‘ show the number of up-regulated genes in each studied comparison. The number of genes of interest are highlighted in a red box. **b**. Heatmap depicts the log2(FC) of the top 50 up-regulated genes in the *V. longisporum* VlsPLA_2_+^WT^ strain as compared to VlsPLA_2_ +^ΔCD^ and WT. Adjusted p-value < 0.05, absolute log2 fold change > 2 for up-regulated genes. Analysis was conducted in four biological replicates.

### VlsPLA_2_ suppresses PTI-related hypersensitive response

It is known that pathogens, especially necrotrophs, secrete a plethora of necrosis-inducing effectors^8^. Thus, we investigated whether VlsPLA_2_ could have the same function by transiently expressing VlsPLA_2_ in *N. benthamiana* plants and monitoring the symptoms daily. No necrosis was observed even seven days after Agro-infiltration (data not shown). Moreover, many pathogens rely on an initial biotrophic stage to establish a successful infection by secreting HR suppressing effectors^28^. Hence, the ability of VlsPLA_2_ to suppress HR was investigated in the Cf4/Avr4 complex. The Cf4 is a Pattern Recognized Receptor (PRR) from tomato plants, recognized by the Avr4 effector produced by fungal pathogen *Cladosporium fulvum*, leading to PTI-related HR ^29,30^. Our results showed that in the area where the VlsPLA_2_^+WT^ was previously Agro-infiltrated, a significant reduction of HR was observed, as compared to the areas where the mock inoculation and the empty-vector control were infiltrated (Fig. 4a). In contrast, the enzymatically inactive version (i.e., VlsPLA_2_^ΔCD^), was not able to suppress HR (Fig. 4a). The role of VlsPLA_2_ in suppressing ETI-induced HR caused by *Pseudomonas syringae* pv *tomato* DC3000 was also investigated. This bacterial strain secretes effectors to the host through a Type III secretion system recognized by the *N. benthamiana* R proteins, leading to a strong HR^31^. Our results showed that VlsPLA_2_ was not able to suppress ETI-related HR, indicating that this effector is involved only to PTI responses (Fig 4b).

**Fig. 4.**
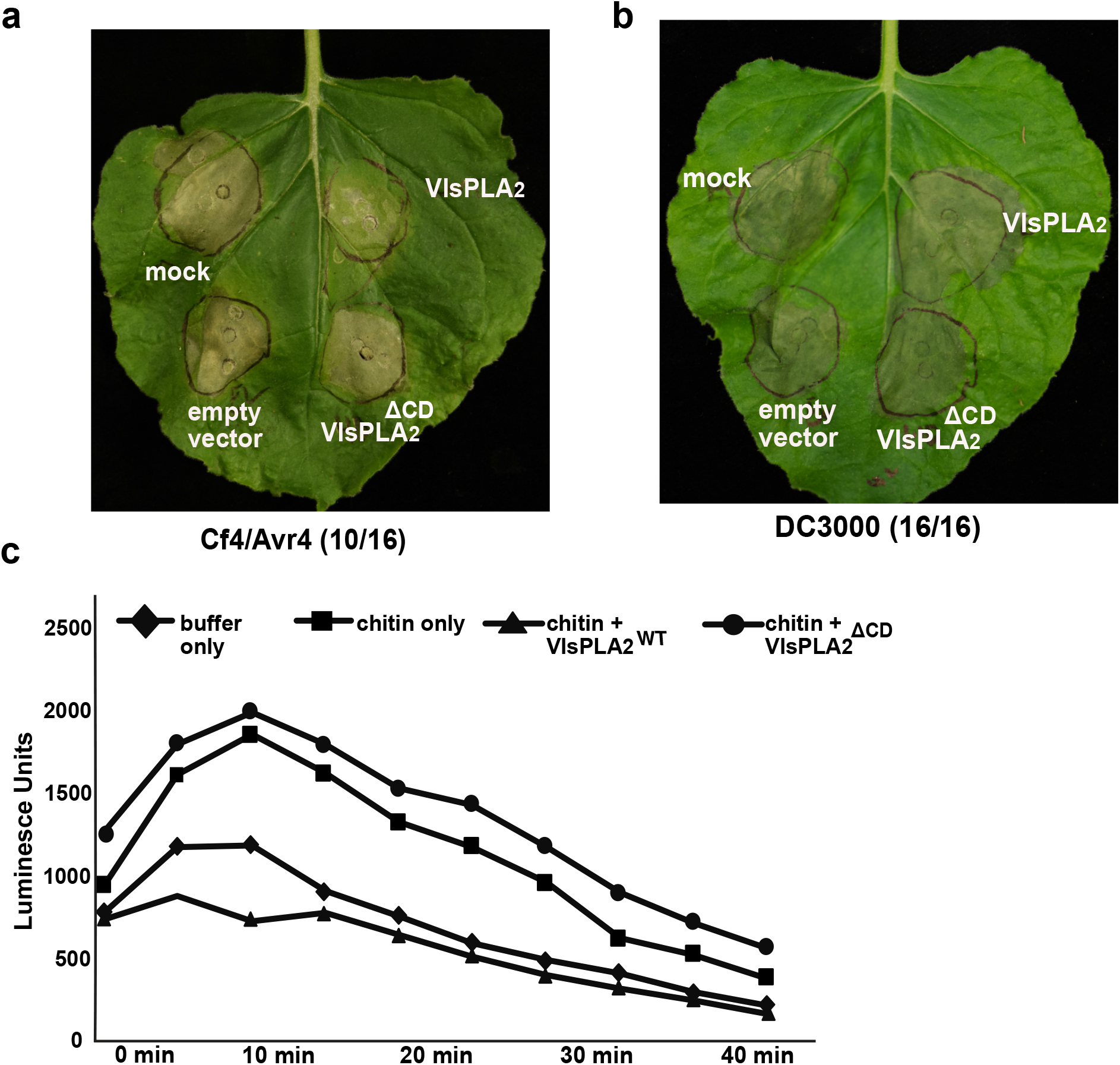
VlsPLA_2_ phospholipase negatively regulates PTI-related hypersensitive responses (HR). **a**. HR suppression assay by VlsPLA_2_^WT^ and VlsPLA_2_^ΔCD^ on *N. benthamiana*, expressing the Cf-4 receptor. HR challenged 24 hours post infiltration with the Avr4 effector. **b** HR suppression assay by VlsPLA_2_^WT^ and VlsPLA_2_^ΔCD^ on *N. benthamiana*. HR was induced by *P. syringae* pv *tomato* DC3000 24 hours post infiltration. Inoculation with the empty pGWB602 vector and mock inoculation with the induction buffer were used as controls. Images taken 3 days post inoculation. In total eight plants and two leaves per plant were used. **b**. Chitin-induced oxidative (ROS) burst assay in *N. benthamiana* leaves. Production of ROS was determined using luminol-dependent chemiluminescence. Leaf discs were treated with chitin, 10 μM VlsPLA_2_^WT^ or VlsPLA_2_^ΔCD^ purified protein, while only buffer was used as a negative control. In total eight biological replicates were used

It is known that ROS signaling plays a key role in the induction of HR^32^. Thus, to investigate whether VlsPLA_2,_ could suppress the ROS burst, a luminol-based protocol was used on *N. benthamiana* leaves. ROS burst was significantly induced by chitin and production was significantly reduced on leaves which previously had been treated with the VlsPLA_2_^+WT^ protein compared to non-treated leaves (Fig 4c). In contrast, no ROS suppression was observed on leaves treated with the enzymatically inactive protein (i.e., VlsPLA_2_^ΔCD^), suggesting that suppression of HR was possibly attributed to the reduction of ROS burst (Fig 4c).

### VlsPLA_2_ is initially localized to the host nuclei

As previously mentioned, the VlsPLA_2_ protein sequences contains two putative NLS motifs (Fig. 1a). Thus, to confirm the predicted subcellular localization of VlsPLA_2_, a construct was made tagging VlsPLA_2_ with GFP at the C-terminus and transiently expressed in *N. benthamiana* leaves, which were monitored using the confocal microscope 48 and 72 hours post infiltration (hpi). Our observations showed that the protein clearly localized to host nuclei at 48hpi, while it was translocated to plant chloroplasts and cell periphery at 72hpi (Fig. 5a). To investigate whether the predicted NLS were functional, a truncated version of VlsPLA_2_ was constructed, where both NLS were disrupted (i.e., VlsPLA_2_ ^ΔNLS1NLS2^). Confocal microscopy showed that this truncated version localized to cell periphery, but no nuclear localization was observed (Fig. 5b). Furthermore, we investigated whether both NLS are critical for nuclear localization and for that reason, truncated versions were constructed where either NLS1 or NLS2 were deleted (i.e., VlsPLA_2_^ΔNLS1^ and VlsPLA_2_^ΔNLS2^). Our analysis showed that both NLS mutants were functional, while the bipartite NLS2 seemed to be more critical for nuclear localization (Supplementary Fig. 4). We also studied whether deletion of the signal peptide (SP) could affect VlsPLA_2_ localization, thus a version lacking this signal was constructed (i.e., VlsPLA_2_^ΔSP^). No changes in nuclear localization were observed between WT and the VlsPLA_2_^ΔSP^ version (Supplementary Fig. 4). We also investigated the localization of the VlsPLA_2_ ^ΔCD^ version at 48hpi. Interestingly, we observed that VlsPLA2^ΔCD^ was localized to the chloroplasts and not able to enter the nucleus, although both NLS were intact, indicating that additional factors could also be involved in its nuclear localization (Fig. 5c).

**Fig. 5.**
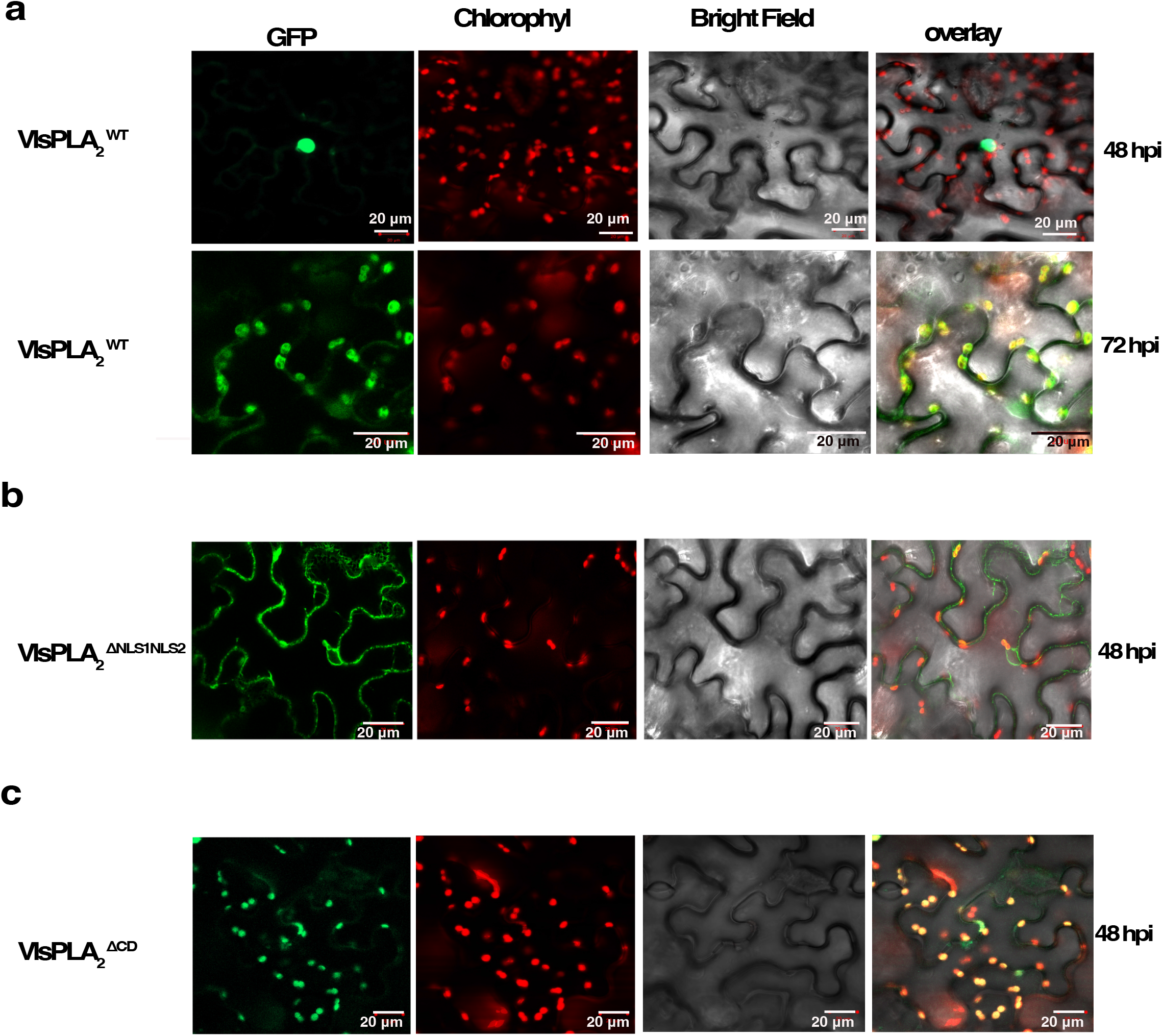
The VlsPLA_2_ phospholipase is initially localized to the host nuclei. Live-cell imaging of **a**. VlsPLA_2_^WT^, **b**. VlsPLA2^ΔNLS1NLS2^ and **c**. VlsPLA_2_^ΔCD^ versions, tagged with GFP at the C-terminus in transiently expressed *N. benthamiana* leaves. The localization was monitored with a laser-scanning confocal microscope with a sequential scanning mode 48 and 72 hours post infiltration. The GFP and the chlorophyll were excited at 488 nm. GFP (green) and chlorophyll (red) fluorescent signals were collected at 505– 525 and 680–700 nm, respectively.

### VlsPLA_2_ binds to the vesicle associated membrane protein A

In order to investigate, whether VlsPLA_2_ potentially interacts with plant proteins, the VlsPLA_2_::GFP-CT was transiently expressed in *N. benthamiana*, pull-downed at 48hpi with GFP-trapped beads, and processed for MS/MS spectrometry. In our analysis, a plethora of potential interactions were identified, as compared to free GFP control (Supplementary Table S4). Fifteen potential interacting proteins were selected for further analysis using yeast-two-hybrid (Y2H) assays and focusing mostly on proteins with a predicted nuclear localization, such as transcription factors, histones, and GTP-binding nuclear Ran proteins (Supplementary Table S4). However, according to our analysis none of these selected proteins were able to interact with the VlsPLA_2_ phospholipase.

Since MS/MS analysis showed that VlsPLA_2_ potentially interacts also with variable membrane vesicle proteins, their interaction was further investigated with Y2H assays (Supplementary Table S4). Our results showed that the yeast strains co-transformed with VlsPLA_2_ and NbS00011956g0004 protein, which is putatively a vesicle-associated membrane protein A (VAMPA), denominated as NbVAMPA-1, was able to grow on auxotrophic selection plates (-His, -Ade, -Leu, -Trp), suggesting physical interactions between VlsPLA_2_ and NbVAMPA-1 (Fig. 6a). Further, disruption of NLS seemed not to have any impact in this interaction (Fig. 6a). However, the co-IP assays showed that NbVAMPA-1 interacted only with the VlsPLA_2_^ΔNLS1NLS2^ version (Fig. 6b). This possibly is attributed to the localization and amount of the VlsPLA_2_^ΔNLS1NLS2^ protein as compared to the WT version of VlsPLA_2_. Confocal microscopy confirmed that NbVAMPA-1 was localized to the cell periphery, and co-localization with VlsPLA_2_ in the plant nuclear membrane was also observed (Fig. 6c; Supplementary Fig. 5a).

**Fig. 6.**
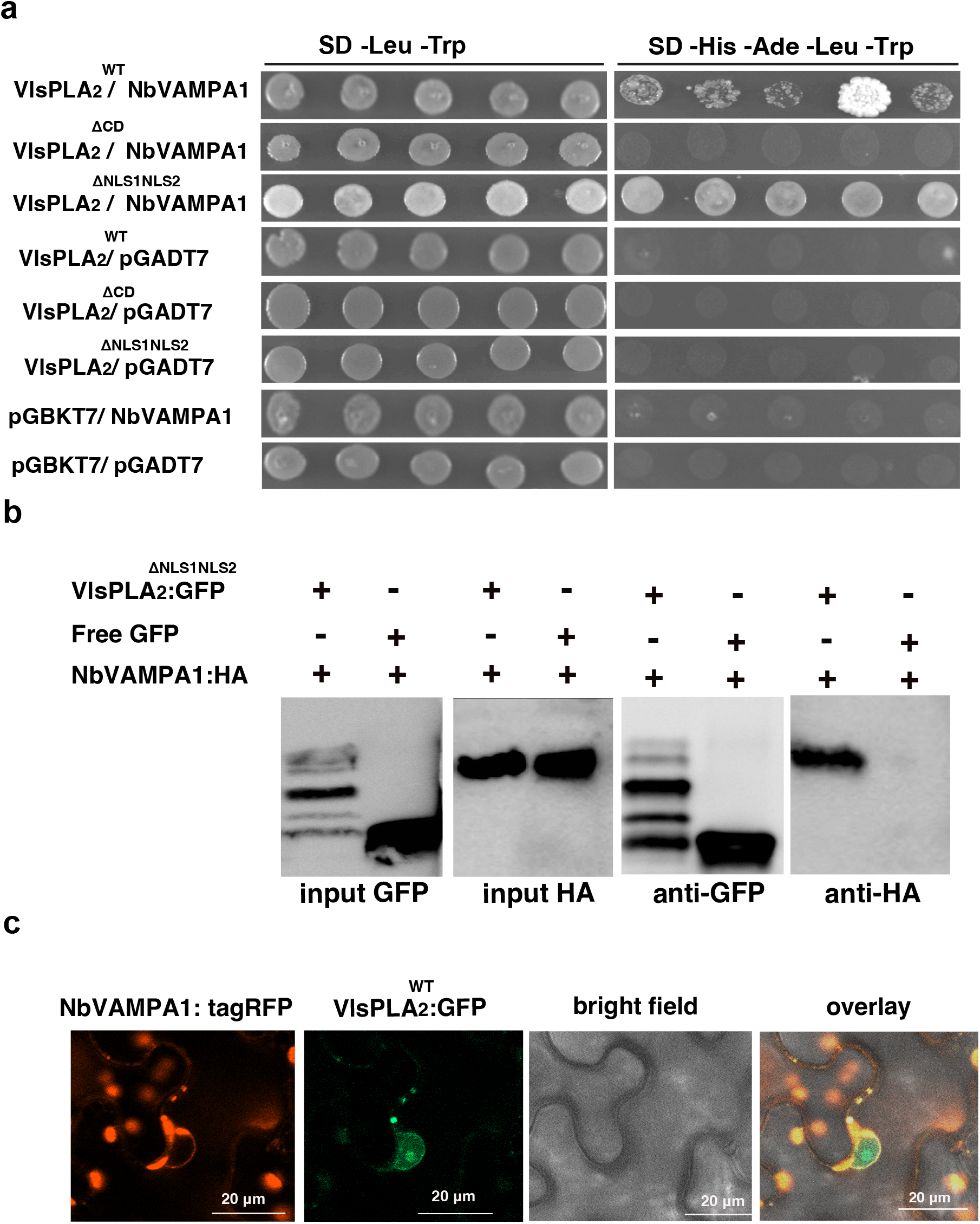
The VlsPLA_2_ phospholipase interacts with the plant vesicle protein NbVAMPA1. **a**. Pairwise yeast-two-hybrid assays between VlsPLA_2_^WT^, VlsPLA_2ΔCD_ or VlsPLA2^ΔNLS1NLS2^ (used as a bait in pGBKT7 vector), and NbVAMPA1 protein (used as a prey in pGADT7 vector). Growth of yeast cells on SD-4 (-His,-Ade, -Leu, -Trp) selective media represents protein–protein interaction and growth on SD-2 (-Leu, -Trp) media confirms yeast transformation. Yeast transformed with the empty vectors were used as negative controls. **b**. Co-immunoprecipitation assay (co-IP) assay between the GFP-tagged VlsPLA_2_^ΔNLS1NLS2^ and HA-tagged NbVAMPA1, transiently co-expressed in *N. benthamiana* leaves and pull downed using the GFP-trap agarose magnetic beads. **c**. Live-cell imaging of VlsPLA_2_^WT^ tagged with GFP at the C-terminus, and NbVAMPA1 tagged with tagRFP at the N-terminus, in Agro-infiltrated *N. benthamiana* leaves. The localization was monitored with a laser-scanning confocal microscope with a sequential scanning mode 48 hrs post infiltration. The GFP and the chlorophyll were excited at 488 nm. GFP (green) and chlorophyll (red) fluorescent signals were collected at 505– 525 and 680–700 nm, respectively. The tagRFP was excited at 558 nm and collected at 545-620 nm.

In addition, we investigated whether the VlsPLA2^ΔCD^ was able to interact with the NbVAMPA-1 protein and interestingly, our results showed no interaction between them (Fig. 6a). Further, prediction of the VlsPLA2^ΔCD^ 3D structure, showed that this mutation did not cause any major change in the protein structure, suggesting that the catalytic residue HD was responsible for the interaction with this vesicle protein.

The NbVAMPA-1 protein structure analysis revealed the presence of a motile sperm protein domain (MSP), which is an ER-anchored receptor involved in interorganelle contacts^33^, at the N-terminus and a transmembrane domain (TM) at the C-terminus. Furthermore, genome analysis of *N. benthamiana* revealed the presence of ten genes that were homologs to the NbVAMPA-1 one. Alignment of these ten homologs showed the presence of a conserved motif of nine amino acids (DMQCKDKFL) (Supplementary Fig. 5b). Thus, to investigate the role of the different domains (MSP, TM) mutants were generated (i.e., NbVAMPA1^ΔMSP^ and NbVAMPA1^ΔTM^ respectively). Our results showed that the MSP-domain-only was not able to interact with VlsPLA_2_ (NbVAMPA1^ΔTM^), similar to the truncated NbVAMPA1 where the MSP domain had been deleted (NbVAMPA1^ΔMSP^), indicating that the intact vesicle protein is required in order to interact with the VlsPLA_2_ phospholipase (Supplementary Fig. 5c). Moreover, the importance of the DMQCKDKFL conserved motif in interactions with VlsPLA_2_ was also investigated. Hence, this conserved motif was mutated (NbVAMPA1^ΔCM^), and protein interactions were evaluated. Our results showed no interaction between VlsPLA_2_ and the NbVAMPA1^ΔCM^ version, suggesting that this conserved motif is important for its binding affinity to VlsPLA_2_ (Supplementary Fig. 5c).

### *VlsPLA*_*2*_ modulates expression of transcription factors and genes involved in plant immunity

To investigate whether the presence of VlsPLA_2_ could cause any modification in the plant’s gene-expression patterns, transcriptome of *N. benthamiana* leaves, transiently expressed *VlsPLA*_*2*_^*WT*^ or *VlsPLA2*^*ΔCD*^, was analysed 48 and 72hpi. *N. benthamiana* leaves transiently expressed the empty vector was used as a control. At 48hpi, 1846 genes were differentially expressed (DEGs) in leaves where the *VlsPLA*_*2*_^*WT*^ gene was expressed, as compared to 144 genes in plants expressing the *VlsPLA*_*2*_ ^*ΔCD*^ (Fig. 7a, Supplementary Table S5). Among these DEGs, 788 were up-regulated, and 1062 were down-regulated (Fig. 7a, Supplementary Table S5). Interestingly, the number of DEGs significantly reduced to 168 at 72hpi; among them, 127 were induced and 52 were suppressed (Fig. 7a, Supplementary Table S5).

**Fig. 7.**
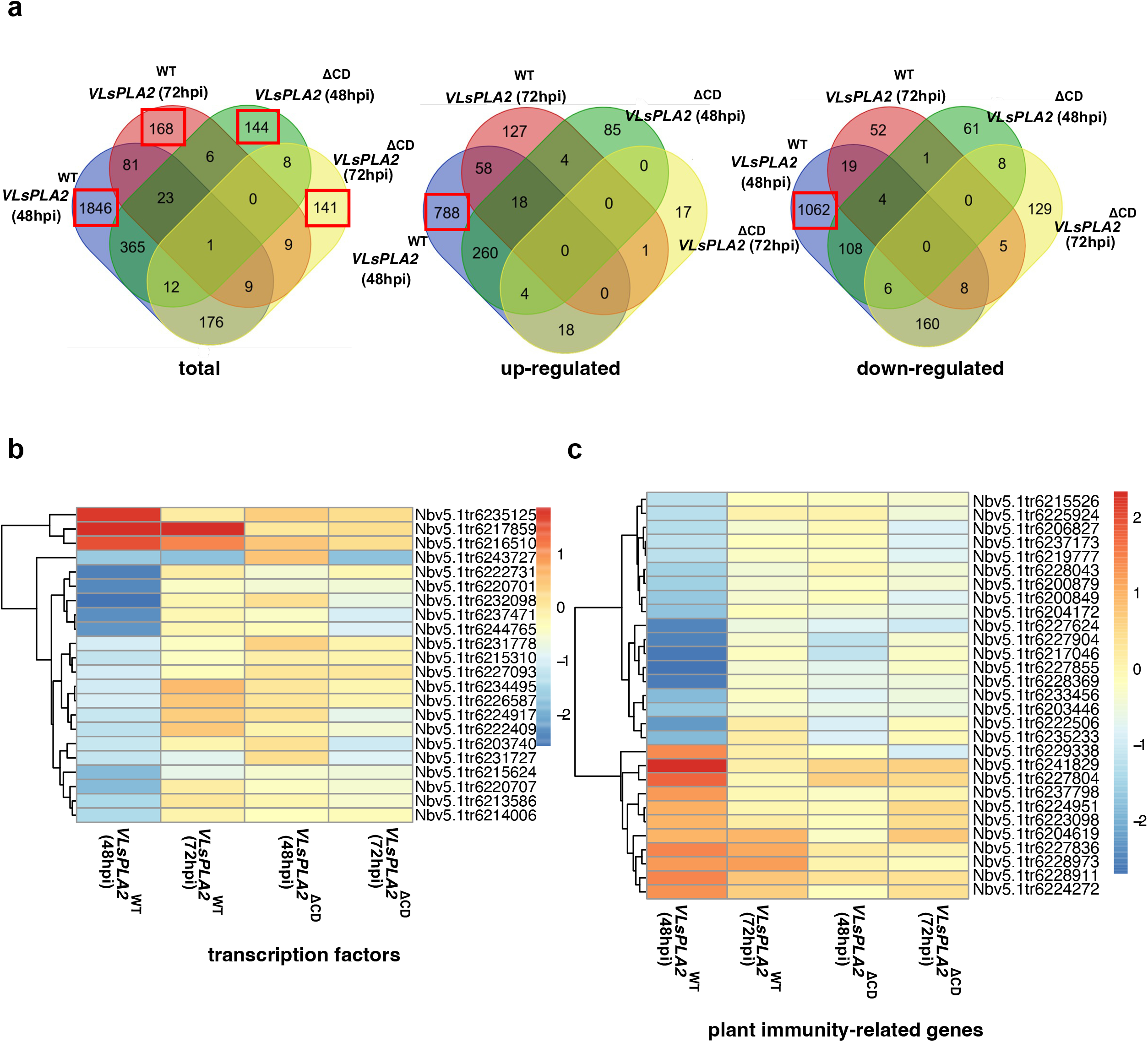
The *VlsPLA*_*2*_ phospholipase modulates plant immunity responses. **a**. Venn diagrams show the number of differentially expressed genes in *N. benthamiana* plants transiently expressing either the functionally active (VlsPLA_2_^WT^) or the inactive (VlsPLA_2_^ΔCD^) VlsPLA phospholipase, 48 and 72 hours post infiltration. The number of genes of interest are highlighted in a red box. **b**. Heatmap depicts the log2(FC) of differentially regulated genes putatively encoding transcription factors in *N. benthamiana* plants transiently expressing either VlsPLA_2_^WT^ or the VlsPLA_2_^ΔCD^ 48 and 72 hours post infiltration. **c**. Heatmap depicts the log2(FC) of differentially regulated genes putatively encoding plant immunity-related proteins in *N. benthamiana* transiently expressing either VlsPLA_2_^WT^ or VlsPLA2^ΔCD^, 48 and 72 hours post infiltration. In all cases the transcriptome of plants inoculated with the empty vector used as a control. (adjusted p-value < 0.05, log2 fold change > 1 for up-regulated genes and < -1 for the downregulated ones). Analysis was conducted in four biological replicates.

Since VlsPLA_2_ initially targets the plant nucleus, we investigated the transcription patterns of genes putatively encoding transcription factors. At 48hpi, in plants where the *VlsPLA*_*2*_^*WT*^ was expressed, we found 22 DEGs encoding transcription factors, as compared to plants expressing the *VlsPLA*_*2*_^*ΔCD*^ or the empty vector (Fig. 7b, Supplementary Table S5). Among them, three were induced and 19 suppressed (Fig. 7b, Supplementary Table S5). Within the down-regulated ones, we found four genes encoding ethylene-responsive transcription factors, four genes encoding basic helix-loop-helix (bHLH) transcription factors, and five genes encoding myeloblastosis (MYB) transcription factors (Fig. 7b, Supplementary Table S5). Moreover, we studied whether the VlsPLA_2_ phospholipase could modulate genes involved in plant immunity. At 48hpi, our data showed that six genes encoding putative receptor-like protein kinases (RLKs) and five genes encoding for Avr9/Cf-9 rapidly elicited proteins (ACRE), were highly induced on plants expressing *VlsPLA*_*2WT*_, compared to plants where the *VlsPLA2*^*ΔCD*^ or the empty vector were expressed (Fig. 7c, Supplementary Table S5). On the other hand, *VlsPLA*_*2*_^*WT*^ expression led to the suppression of 14 genes encoding putatively subtilisin-like proteases and F-box proteins, and three genes encoding aquaporins (Fig. 7c, Supplementary Table S5). In summary, the data from this analysis indicated that VlsPLA_2_ was a strong modulator of plant immunity responses.

## Discussion

In the current study, we investigated the functional role of a fungal secreted PLA_2_ enzyme, with emphasis on its involvement in infection biology of the phytopathogen *V. longisporum*. Identification of PLA_2_ genes only in 28 fungal genomes showed patchy distribution of this gene across the fungal kingdom, with no clear correlation to ecological niches or lifestyles. This result is in line with the previous finding, which suggested that *PLA*_*2*_ probably derived through horizontal gene transfer from a prokaryotic ancestor to *V. longisporum*^*34*^. The expression of *VlsPLA*_*2*_ was induced in *V. longisporum* during interaction with *B. napus* roots. This is consistent with the previous findings where induction of a gene encoding a secreted *PLA*_*2*_ was observed during the establishment of symbiosis in the mycorrhizal species, *Tuber borchii*. It was also activated during autoproteolytic activity and promoted mycorrhiza development with a specific host plant^23,35,36^, suggesting that secreted PLA_2_ could play an important role in different aspects of fungal-host interactions.

Previous studies regarding the role of secreted PLA_2_ in virulence have been conducted mainly in bacterial-mammalian cell interactions, in which they have been shown to work as toxins inducing proteolysis and hemolysis (e.g., ExoU produced by *Pseudomonas aeruginosa* and SlaA produced by Group A *Streptococcus*), and thereby increasing disease severity^37,38^. It has been proposed that secreted bacterial PLA_2_ could degrade the phospholipid components in the host plasma membranes, releasing arachidonic acid (AA) and the subsequent synthesis of eicosanoids, which interferes in signal transduction cascades, altering the transcription of genes and leading to increased inflammatory responses^6^. A similar function could be speculated for the VlsPLA_2_ phospholipase during interactions with the plant host. Our data from this study showed that that overexpression of *VlsPLA*_*2*_ in fungal cells resulted in increased *V. longisporum* virulence, and heterologous expression of this phospholipase in *A. thaliana* led to an increased susceptibility against this pathogen. To establish a mechanistic connection between *VlsPLA*_*2*_ overexpression and increased virulence, transcriptome of both *V. longisporum* mycelia and *N. benthamiana* leaves, overexpressing *VlsPLA*_*2*_, were analyzed. Induced expression of a diverse range of virulence factor genes, encoding zinc finger proteins, cytochrome monooxygenases, efflux-pump proteins, and a Calpain-A like protein^25,26,27^ was observed, indicating that this phospholipase is involved in signal transduction cascades in fungal cells, altering the expression of genes and leading to increased virulence. Similarly, the transcriptome analysis of *N. benthamiana* leaves showed that expression of VlsPLA_2_ was able to induce RLKs and ACREs, which encode signalling pathways involved in the initial stages of defence responses^39^. We also observed that expression of this phospholipase suppressed many genes encoding MYB and bHLH transcription factors in the plant. These types of transcription factors have been shown to be involved in the regulation of plant defence mechanisms, such as jasmonic acid (JA)-mediated and HR-triggered responses^40,^. In addition, expression of VlsPLA_2_ led to suppression of a number of genes encoding subtilisin-like proteases and F-box proteins. Their role in the activation of downstream immune responses, including involvement in PTI-related HR, has already been shown^41,42,43^ Moreover, these data were supported by the result that VlsPLA_2_ was able to suppress PTI-related HR, but not the ETI-induced one, indicating that this phospholipase plays a role only in suppression of basal immune responses. Taken together, these data indicate that VlsPLA_2_ is a strong modulator of basal plant immune responses.

The nuclear localization of PLA_2_ was previously observed in human endothelial cells, and in *Arabidopsis thaliana* plants^44,45^. Especially in *A. thaliana*, PLA_2_ suppresses cell death^45^, similar to the VlsPLA_2_ one. It has previously been showed that the nuclear translocation of a cytosolic PLA_2_ elicits localized hydrolysis of phospholipids in rat cells^46^. This agrees with our data, showing that VlsPLA_2_ increased the production of certain phospholipids; thus, we could suggest that the products from this enzymatic function are possibly involved in signal transduction pathways and altering the expression of genes involved in plant immunity. We also found that the VlsPLA_2_ phospholipase was associated with a plant VAMP-A protein. These are highly conserved proteins present in all eukaryotes and are involved in several physiological functions including membrane trafficking, lipid transport, and unfolded protein response, among others^47^. It has been shown that VAMPs could interact with many different proteins, including proteins involved in plant resistance against fungal pathogens^48,49,50^. Our data showed that NbVAMPA1 promotes the entry of VlsPLA_2_ to the plant nucleoplasmic reticulum and, presumably, this membrane protein is binding to the active site of VlsPLA_2_. Similar results have observed in the mammalian cells, where a VAMP-A protein and an oxysterol-binding protein promote the entry of endosomes into the nuclear envelope^51^.

In conclusion, the role of a secreted VlsPA_2_ enzyme in the virulence of the phytopathogenic fungus *V. longisporum* has been investigated. This phospholipase is a regulator of fungal genes, involved in virulence and targets the plant nuclei by hijacking the VAMP membrane proteins. Once there, VlsPA_2_ suppresses the expression of genes that play a crucial role in induction of basal immune responses, possibly through the production of phospholipids and their involvement in signal transduction cascades. To the best of our knowledge, this study is the first to examine in detail a fungal secreted PLA_2_, by giving important insights about its role in fungal infection biology.

## Material and Methods

### Fungal strains and growth conditions

In the current study, the *V. longisporum* isolate VL1 (CBS110220) was used. The fungus was kept on potato dextrose agar (PDA, Difco) at 20°C in darkness and sporulation occurred within seven days on this medium.

### Nucleic acids manipulation and gene expression analysis

DNA was extracted using the NucleoSpin Plant II kit (Macherey-Nagel GmbH), while RNA was extracted using the Spectrum Plant Total RNA kit (Sigma). RNA was reverse transcribed using the iScript cDNA synthesis kit (Bio-Rad) and the qPCR analysis was done using the SsoFast EvaGreen supermix (Bio-Rad). All PCR reactions were done using Phusion Hot Start II High-Fidelity PCR Master Mix (Thermo Scientific). All PCR products were gel-purified using the GeneJET Gel Extraction Kit (Thermo Scientific). One Shot® TOP10 Chemically Competent *E. coli* (Invitrogen) was used for routine cloning and plasmids were isolated using the GeneJET Plasmid Miniprep Kit (Thermo Scientific). All the kits were used following the manufacturers’ protocol.

The transcription profiles of *V. longisporum* genes encoding candidate effector proteins were investigated under conditions previously described^52^. Briefly, *B. napus* seedlings cultivar ‘‘Hannah’’ were inoculated with *V. longisporum* by dipping in 10^6^ conidia/ml suspension for 15min. Plants were harvested at 2-, 4-, 6-, 8- and 10-dpi and frozen in liquid nitrogen, while mycelia grown in potato dextrose broth (PDB, Difco) were used as a control. RNA was extracted and treated with DNase I (Thermo Fisher Scientific). For RT-qPCR, one μg of total RNA was reverse transcribed and used for the analysis. Primers sequences are listed in Supplementary Table S6, and gene expression was normalized using the reference gene, glyceraldehyde phosphate dehydrogenase (*GAPDH*)^53^. Relative expression values were calculated from the threshold cycle (Ct) values according to the 2^−ΔΔCT^ method^54^.

### Protein sequence analysis, structure prediction, and phylogeny

The amino acid sequence of VlsPLA_2_ was retrieved from the *V. longisporum* VL1 genome^21^. Analysis of conserved domains was performed using the SMART protein tools^55^. To investigate whether VlsPLA_2_ is an apoplastic or a cytoplasmic effector, the ApoplastP prediction tool was used, and its subcellular localization was predicted using the LOCALIZER software^56,57^.

Fungal genomes retrieved from the MycoCosm database and searched for putative secreted *PLA*_*2*_ genes^58^. For the phylogenetic analysis, homologs to VlsPLA_2_ were retrieved and the amino acid sequences were aligned with CLUSTAL W and analysis was implemented in MEGA X, using the JTT amino acid substitution model^59^. Statistical support for branches was supported by 1000 bootstraps. Finally, prediction of the VlsPLA_2_ 3D structure was conducted using the SWISS-MODEL server^60^.

### Plasmids and vectors generated in the study

Gene fragments for VlsPLA_2_ and NbVAMPA1 WT and truncated versions, were amplified from *V. longisporum* or *N. benthamiana* cDNA, respectively, or synthesized and inserted in the pUC19 plasmid by Integrated DNA Technologies (Coralville, IA). They were subcloned into the entry vector pDONOR/Zeo (Invitrogen) using Gateway™ BP Clonase™ enzyme mix (Invitrogen), by following the manufacturer’s protocol. The different entry vectors were cloned with the Gateway™ LR Clonase™ enzyme mix (Invitrogen), to the destination vectors, showed in Supplementary Table S7. The expression of genes *in planta*, were driven by the *35S::CaMV* constitutive promoter.

To overexpress VlsPLA2+^WT^ or VlsPLA2+^ΔCD^ in the fungus, the pRFHUE vector was used, which contains the *gpdA* constitutively expressed promoter. Briefly, the wild type phospholipase gene (VlsPLA_2_^WT^) was amplified from *V. longisporum* cDNA, while the VlsPLA_2_^ΔCD^ gene was amplified from the plasmid pUC19-VlsPLA_2_^ΔCD^. The genes and the backbone of the vector were amplified using the primers enlisted in the Supplementary Table S6. The PCR products were gel purified and ligated using the GeneArt™ Seamless Cloning and Assembly Enzyme Mix (Invitrogen), following the manufacturer’s protocol.

### Heterologous protein expression and enzymatic activity assay

The subcloning, expression, and purification of the active (VlsPLA_2_^WT^) and inactive (VlsPLA2^ΔCD^) protein was done at the Protein Expertise Platform (PEP) at Umeå University. Briefly, Origami2 (DE3) cells were grown in LB media supplemented with 50 μg/mL kanamycin at 37°C with agitation. The culture was incubated until an OD_600_ ∼ 0.6–0.7 was achieved. Protein expression was induced by the addition of 0.4 mM isopropyl β-D-1-thiogalactopyranoside (IPTG) and incubated overnight at 20°C. Cells were harvested by centrifugation 4000 x g for 20min. The target protein was purified using His-Tag Purification Resin (Roche) according to the manufacturer instructions.

The His-Trx-tag was cleaved off using Prescission3C protease. Finally, the protein was run on a Superdex200 (Cytiva), 20mM NaP 8.0, 200mM NaCl, 0.5mM b-merchaptethanol. For the phospholipase A_2_ enzymatic activity, the EnzChek Phospholipase A_2_ activity kit was used (Thermo Fisher), according to manufacturer instructions. In all assays, 10μM of pure active (VlsPLA_2_^WT^) and inactive (VlsPLA2^ΔCD^) protein was used. PLA_2_ from *Apis melifera* venom was used as a positive control, and enzymatic activity was monitored at 515 nm using the BMG LABTECH microplate reader.

### Lipidomic analysis

For the lipidomic analysis, overnight cultures of *Agrobacterium tumefaciens* (C58C1) harboring either the pGWB602-VlsPLA_2_^WT^; pGWB602-VlsPLA_2_^ΔCD^; or the empty vector (i.e., pGWB602) were infiltrated on leaves of 4-week-old *Nicotiana benthamiana* plants (grown under 18h light/6h dark at 23°C). Leaves were harvested at 48hpi and lipids were extracted using the chloroform:methanol (2:1) phase extraction method. The chloroform phase was used for UHPLC-QTOF/MS analysis, using the 1290 Infinity UHPLC (Agilent), coupled to the QTOF 6546 instrument (ODIN) (Agilent), using the parameters previously described for this type of analysis^61^. Data analysis was conducted using the ProFinder 10.0 software (Agilent), by comparing the peaks with the internal database standards for different lipid classes.

### Fungal transformation and virulence assays

The vectors pRFHUE-VlsPLA_2_^WT^ and pRFHUE-VlsPLA_2_^ΔCD^ were transformed into *V. longisporum* using the *Agrobacterium*-mediated protocol^62^. The selection of transformed fungal strains was accomplished using 50 μg/mL of hygromycin. The expression levels of the *VlsPLA*_*2*_ gene in the mutant strains was investigated by RT-qPCR techniques as described above. Single-spore cultures were performed from the five colonies with the highest expression and used for the virulence assays. The growth rate of *V. longisporum* overexpression strains were evaluated on PDA plates. For the virulence assays, the plant species *Arabidopsis thaliana* was used. Seeds were surface sterilized and grown in vitro on half strength Murashige and Skoog medium (½MS; Duchefa Biochemie) for two weeks. Plant infections with *V. longisporum* WT and the overexpression strains (i.e., *VlsPLA*_*2*_^WT^ and *VlsPLA*_*2*_^ΔCD^), as well as mocked-inoculated ones, was conducted as described above. Then, plants were transferred to peat soil and grown in short-day conditions (8 hours light, 16 hours dark) at 23°C/18°C for four weeks and watered regularly. The severity of disease was categorized as healthy; mildly infected; severely infected; and dead plants. In addition, the rosette diameter was measured at 28 days post infection.

### Construction *of Arabidopsis thaliana* transgenic plants

*Agrobacterium tumefaciens* strain C58C1 cells carrying the vector pGWB605-VlsPLA_2_^WT^, were used to transform *A. thaliana* Col-0 using the floral dip method as and 50 μg/ml of BASTA (glufosinate ammonium) to select for transformants. Expression levels of VlsPLA_2_^WT^ in the transgenic lines was confirmed by RT-PCR and Western blot. Two independent homogenous lines in T4 generation were used for the virulence assays, as described above.

### Live-cell imaging, hypersensitive response, and ROS burst suppression assays

Overnight cultures of *A. tumefaciens* (C58C1) harboring the vectors, showed in Supplementary Table S7, were infiltrated on leaves of 4-week-old *N. benthamiana* plants (grown under 18h light/6h dark at 23°C), The subcellular localization of the phospholipase was monitored using a Zeiss LSM 800 confocal microscope at 48 and 72hpi.

For the HR suppression assay, the pGWB602-VlsPLA_2_^WT^ and pGWB602-VlsPLA_2_ ^ΔCD^ vectors were transiently expressed by in *N. benthamiana* plants harboring the Cf-4 receptor protein from tomato plants. The HR was triggered 24hrs after the Agro-infiltration with the *Cladosporium fulvum* Avr4 effector protein at OD_600_ = 0.03. The ability of VlsPLA_2_ to suppress ETI-triggered HR was investigated on *N. benthamiana* wild type plants upon infection with *Pseudomonas syringae* pv. *tomato* DC3000. Agro-infiltration with empty vector and only induction buffer (mock), were used as controls. For suppression of chitin-induced oxidative burst of reactive oxygen species (ROS), a luminol-based protocol was used. Briefly, leaf discs from *N. benthamiana* plants, treated with chitin (100 μl/ml), luminol (200μM) (Sigma) and 10 μg/ml horseradish peroxidase (Sigma). The suppression of ROS was analyzed by using 10 μM VlsPLA_2_^WT^ and VlsPLA2_2_^ΔCD^ proteins and measuring the chemiluminescence levels in the BMG LABTECH microplate reader.

### MS/MS spectrometry assay

The pGWB605-VlsPLA_2_^WT^ vector, was transiently expressed in *N. benthamiana*, while free GFP vector was used as a negative control. Proteins were extracted using an extraction buffer containing 20 mM HEPES pH 6.8, 150 mM NaCl, 1 mM EDTA, 1 mM DTT, 0.5% Tween 20, 1mM PMFS and proteases inhibitor cocktail (Roche) and pull-downed using the GFP-trap agarose magnetic beads (Chromotek). Samples proceeded for LC-ESI-MS/MS analysis at the Clinical Proteomics Mass Spectrometry facility, Karolinska Institute/ Karolinska University Hospital/ Science for Life Laboratory, Stockholm. On-bead reduction, alkylation, and digestion (trypsin, sequencing grade modified, Pierce) was performed followed by SP3 peptide clean-up of the resulting supernatant. Each sample was separated using a Thermo Scientific Dionex nano LC-system in a 4 hr 5-40 % ACN gradient coupled to Thermo Scientific High Field QExactive. The software Proteome Discoverer vs. 1.4, including Sequest-Percolator for improved identification, was used to search the *N. benthamiana* v044 database for protein identification, limited to a false discovery rate of 1%.

### Co-immunoprecipitation and yeast-two-hybrid assays

For the yeast-two-hybrid assays, vectors were transformed into the *Saccharomyces cerevisiae* AH109 strain (Clontech). Transformations with empty vectors were used as negative controls. For co-immunoprecipitation assays, pGWB605-VlsPLA_2_^WT^ or pGWB605-VlsPLA_2_ ^ΔNLS1NLS2^, transiently co-expressed with pGWB614-NbVAMPA1 in *N. benthamiana* and pull-downed as described above for the MS/MS assays. GFP-tagged protein was detected using the B2 anti-GFP HRP-conjugated antibody (Santa Cruz Biotechnology), and HA-tagged proteases were detected using the anti-HA peroxidase-conjugated antibody (Sigma), according to manufacturers’ instructions.

### RNA sequencing and bioinformatics analysis

For the transcriptomic analysis in *V. longisporum* mycelia, the WT, *VlsPLA2+*^WT^ and *VlsPLA*_*2*_*+*^ΔCD^ strains were grown in PDB (Difco) for five days at 20°C. For the *in planta* transcriptomic analysis, *N. benthamiana* plants were Agro-infitrated with the pGWB602-VlsPLA_2_^WT^ and pGWB602-VlsPLA_2_^ΔCD^ as well as the empty vector (i.e., pGWB602), as described above. The infiltrated leave areas were harvested 48 and 72hpi for RNA extraction. In both cases, total RNA was extracted with the Plant Total RNA kit (Sigma-Aldrich), and 1000 ng of total RNA was treated with DNase I (Thermo Fisher Scientific), before the generation of RNA strand-specific libraries, using the TruSeq stranded mRNA library preparation kit with polyA selection (Illumina, Inc). Libraries were sequenced using the Illumina NovaSeq 6000 at the SNP&SEQ Technology Platform, Science for Life Laboratory at Uppsala University, Sweden. The experiment was performed in four biological replicates.

The bioinformatics analysis was done as follows. Reads were cleaned and adapters were removed with bbduk v. 38.9^63^ with the following parameters: ktrim=r k=23 mink=11 hdist=1 tpe tbo qtrim=r trimq=10). The quality of the trimmed reads was evaluated using the fastqc v.11.9 quality control tool^64^. *Verticillium longisporum* reads were then mapped to *V. longisporum* VL1 genome (accession GCA_001268145.1) using the splice-aware aligner STAR v. 2.7.9a^65^ with default parameters. The *N. benthamiana* reads were mapped to the *N. benthamiana* v.5.1 transcriptome (http://sefapps02.qut.edu.au/benWeb/subpages/downloads.php) using the non-splice-aware aligner bowtie2 v. 2.4.2.^66^. The number of reads mapping to each gene was quantified using featureCounts v. 2.0.1^67^ considering only sense reads, and differential expression was determined with the DESeq2 R package v. 1.28.1^68^ using a minimal threshold of 1 for log2(FC) and 0.05 for FDR adjusted pvalue. Data visualization was performed with R pheatmap module.

For functional annotation, protein sequences were extracted from the genomes and gene coordinates files of *N. benthamiana* and *V. longisporum* using GffRead v. 0.12.2 ^69^, InterProScan v.5.46.81.0 was used with default parameters to determine additional domains. *V. longisporum* proteins were compared to the PHI-base database v. 09-05-2022^70^ using blastp v.2.13.0+ with minimum 80% in both identity and query coverage to determine known factors of virulence.

## Data availability

The transcriptome data of this study (*V. longisporum* and *N. benthamiana* reads) were deposited on ENA database under Bioproject accession number PRJEB55988.

## Acknowledgments

We would like to thank Wageningen University for providing us with *N. benthamiana* Cf4 seeds, the Protein Expertise Platform (PEP) for the cloning, expression, and purification of VlsPLA_2_, and the Swedish Metabolomics Centre (SMC) for the lipidomic analysis at Umeå University. We would also like to thank the Clinical Proteomics Mass Spectrometry facility, at Karolinska Institute, for MS/MS spectrometry, and the SNP&SEQ Platform at SciLife Lab, Uppsala for RNA-seq analyses. We are also grateful to Anna Tornkvist for helping us with the confocal microscopy, and to Jerry Stålberg for the prediction of the VlsPLA_2_ protein structure and analysis.

---

**Supplementary Fig. 1.**
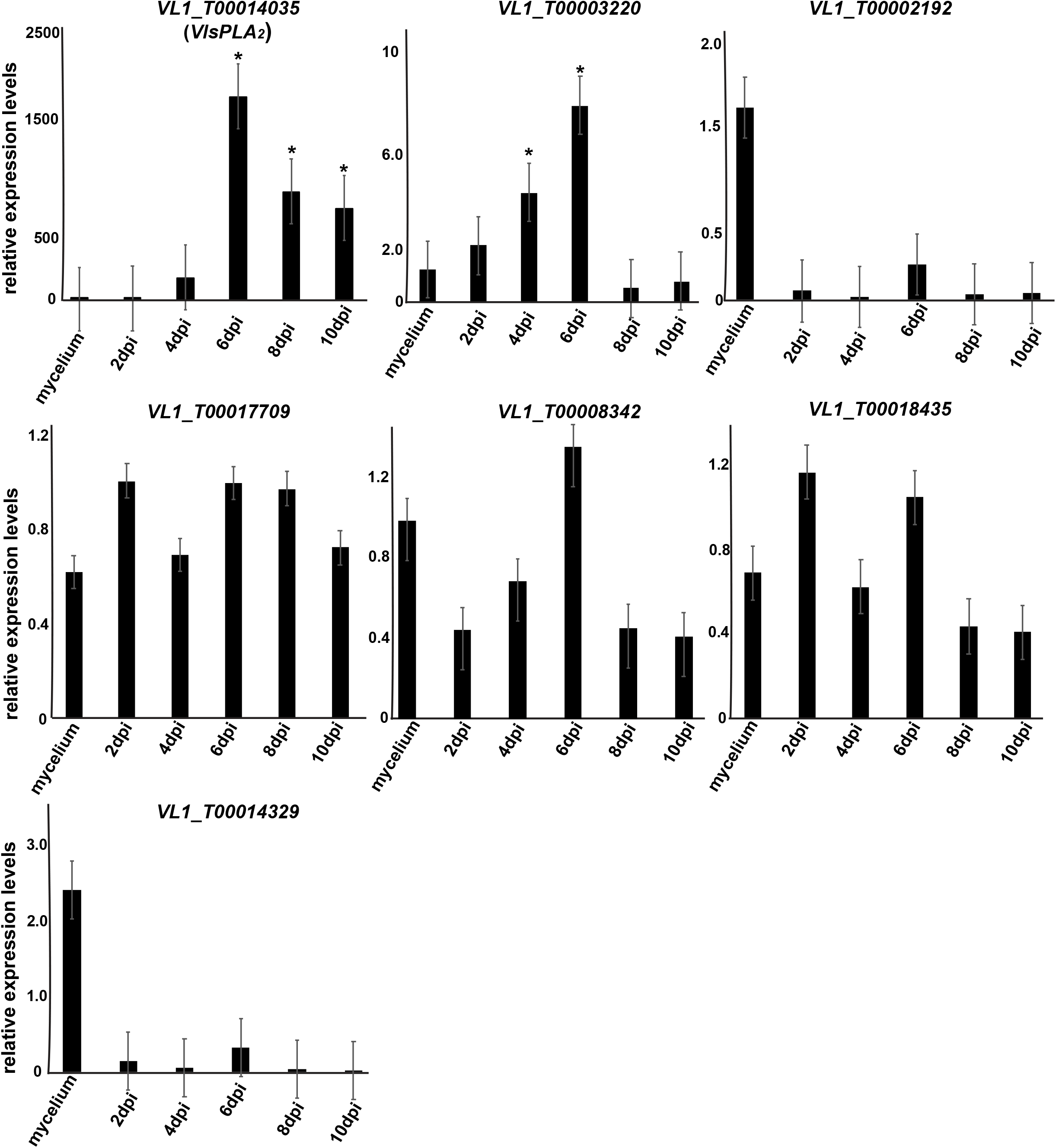
Transcription profiles of *V. longisporum* genes encoding candidate effector proteins upon infection of *Brassica napus*. Gene expression analysis was conducted according to 2^−ΔΔCT^ method. Data were normalized using the expression levels of the reference gene, glyceraldehyde phosphate dehydrogenase (*GAPDH*). Error bars represent SE based on at least three biological replicates. Asterisks indicate statistically significant differences according to Students T’ test (p value < 0.05).

**Supplementary Fig. S2.**
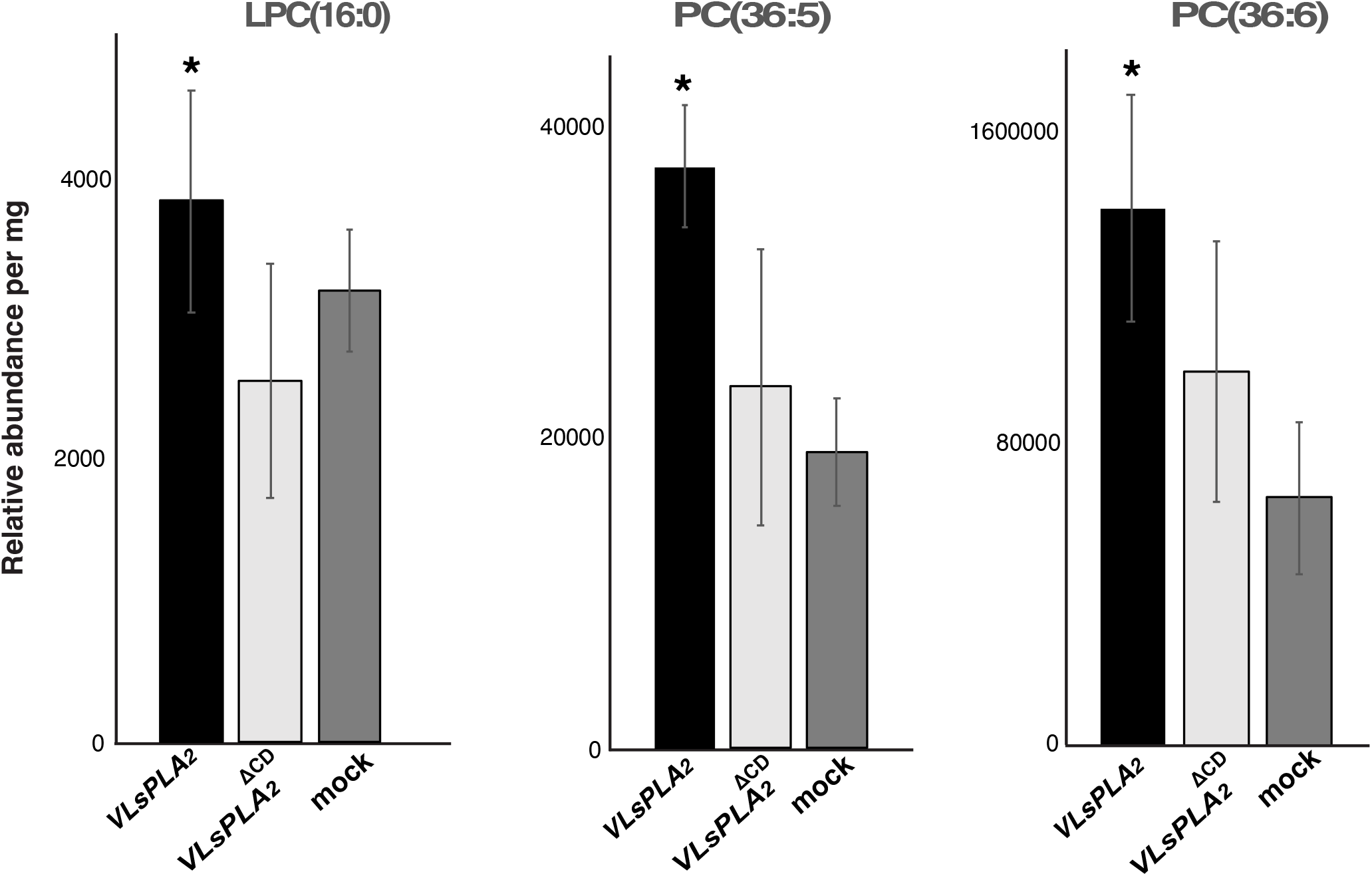
Phospholipid profiles on *N. benthamiana* transiently expressed either VlsPLA_2_ ^WT^ or VlsPLA_2_ ^ΔCD^. Mocked-inoculated plants were used as control. Data depicted as a relative abundance of phospholipids per mg of dry plant tissue. Asterisks (*) indicate statistically significant differences according to the Student’s T test (p < 0.05). Bars represent the standard error (SE) based on five biological replicates.

**Supplementary Fig. S3.**
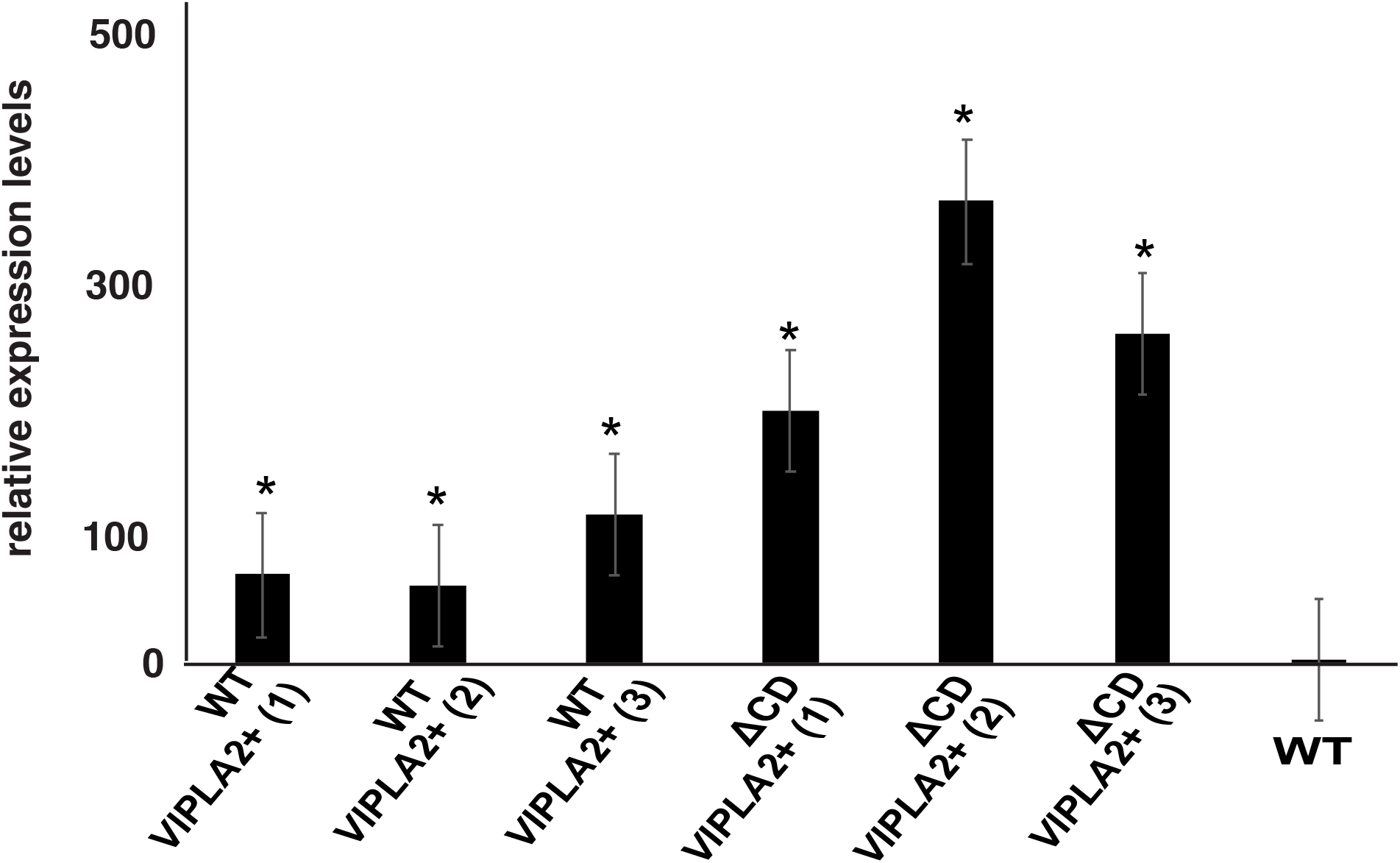
Transcription analysis on *Verticillium longisporum* strains overexpressing either the functional active (VlsPLA_2_+^WT^) or the inactive (VlsPLA_2_ + ^ΔCD^) VlsPLA_2_ phospholipase. Gene expression analysis was conducted according to 2^−ΔΔCT^ method. Data were normalized using the expression levels of the reference gene,glyceraldehyde phosphate dehydrogenase (*GAPDH*). Asterisks indicate statistically significant differences between the overexpression and WT strains according to the Students T’ test (p value < 0.05).

**Supplementary Figure S4.**
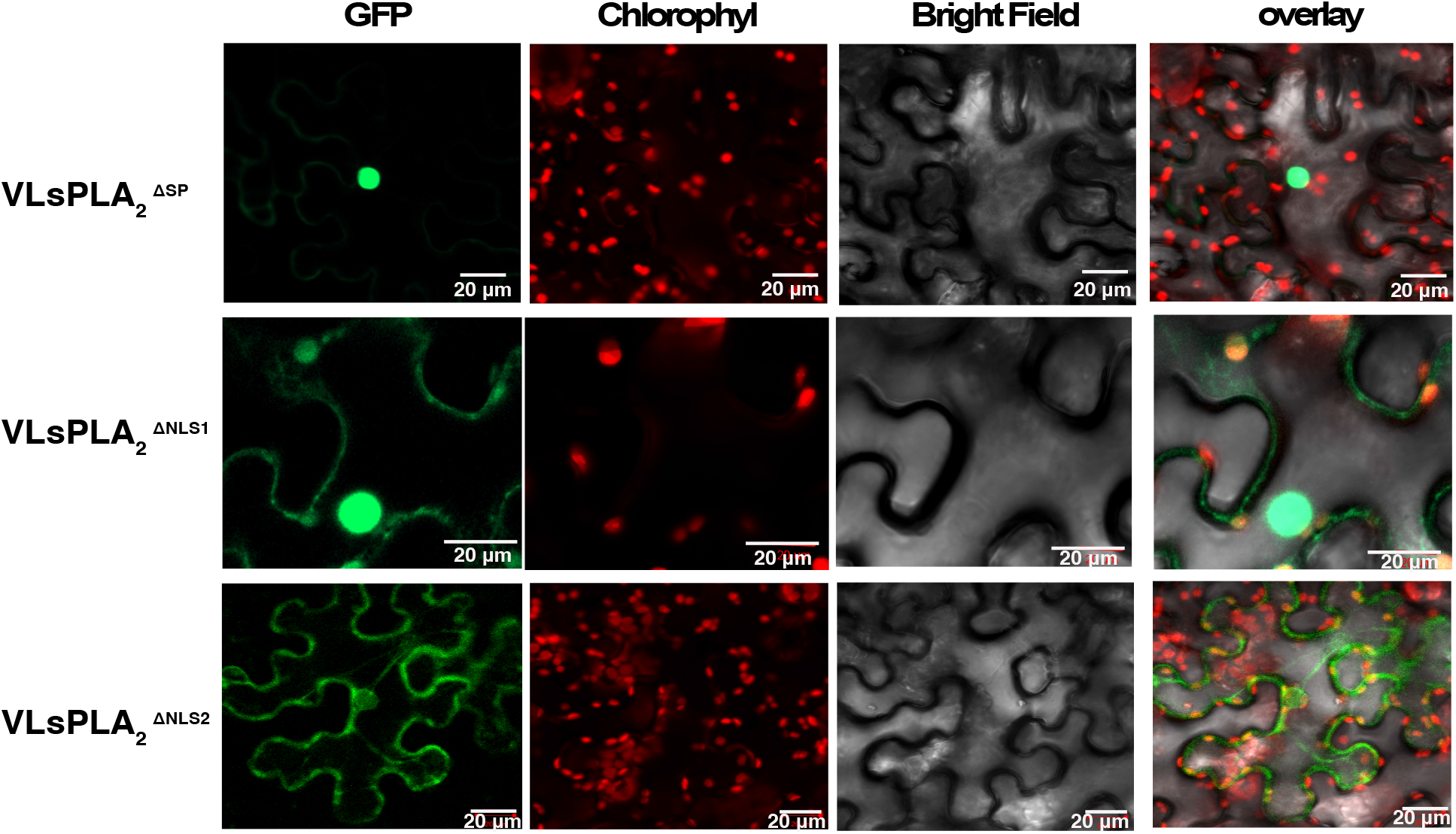
Live-cell imaging of VlsPLA_2_ mutants where signal peptide (VLsPLA_2_^ΔSP^), or NLS1 (VLsPLA_2_ ^ΔNLS1^), or NLS2 (VLsPLA_2_ ^ΔNLS2^) were truncated. Mutants tagged with GFP at the C-terminal and Agro-infiltrated *N. benthamiana* leaves. The localization was monitored with a laser-scanning confocal microscope with a sequential scanning mode 48 hours post infiltration. The GFP and the chlorophyll were excited at 488 nm. GFP (green) and chlorophyll (red) fluorescent signals were collected at 505– 525 and 680–700 nm, respectively.

**Supplementary Fig. S5.**
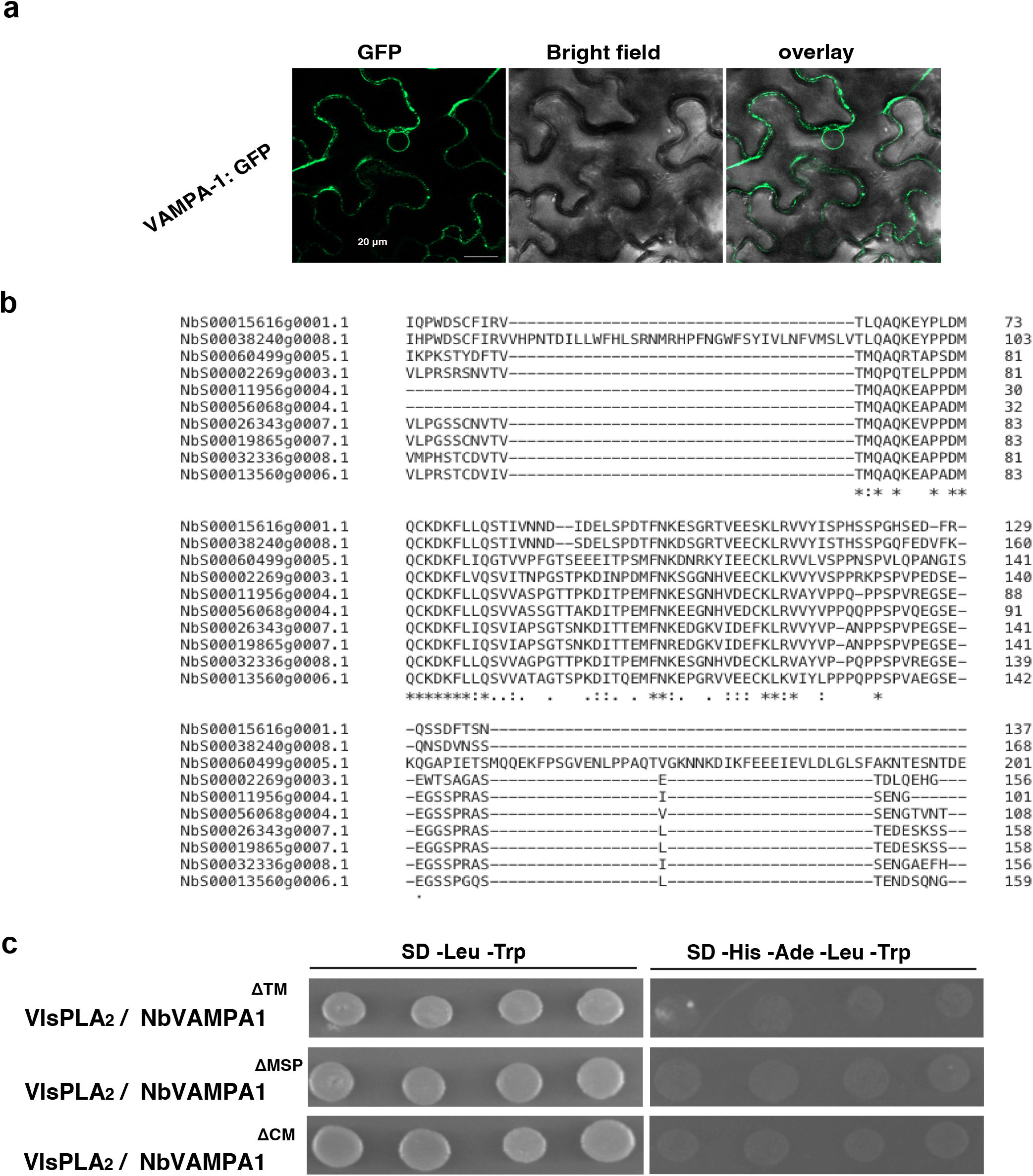
Analysis of NbVAMPA1 protein. **a**. Live-cell imaging of NbVAMPA-1 tagged with GFP at the N-terminal and Agro-infiltrated *N. benthamiana* leaves. The localization was monitored with a laser-scanning confocal microscope with a sequential scanning mode 48 hours post infiltration. The GFP was excited at 488 nm and collected at 505– 525 nm. **b**. Alignment of homologs to NbVAMPA1 in *N. benthamiana*. Identical sequences are marked with asterisks (*). **c**. Pairwise yeast-two-hybrid assays between VlsPLA_2_ (used as a bait in pGBKT7 vector) and NbVAMPA1 mutated versions (used as a prey in pGADT7 vector). Growth of yeast cells on SD-4 (-His, -Ade, -Leu, -Trp) selective media represents protein–protein interaction and growth on SD-2 (-Leu, -Trp) media confirms yeast transformation

## References

1. Nakamura, Y. Plant phospholipid diversity: emerging functions in metabolism and protein-lipid Interactions. Trends in Plant Sci. 22, 1027–1040 (2017).

2. Aloulou, A., Ben Ali, Y., Bezzine, S., Gargouri, Y. & Gelb, M.H. Phospholipases: An Overview. Lipases and Phospholipases: Methods and Protocols 861, 63–85 (2012).

3. Hong, Y.Y. et al. Plant phospholipases D and C and their diverse functions in stress responses. Prog. Lipid Res. 62, 55–74 (2016).

4. Zhao, J. Phospholipase D and phosphatidic acid in plant defence response: from protein-protein and lipid-protein interactions to hormone signalling. J. Exp. Bot. 66, 1721–1736 (2015).

5. Ghannoum, M.A. Potential role of phospholipases in virulence and fungal pathogenesis. Clin. Microbiol. Rev. 13, 122–143 (2000).

6. Sitkiewicz, I., Stockbauer, K.E., & Musser, J.M. Secreted bacterial phospholipase A2 enzymes: better living through phospholipolysis. Trends Microbiol. 15, 63–69 (2007).

7. Giraldo, M.C. & Valent, B.. Filamentous plant pathogen effectors in action. Nat. Rev. Microbiol. 11, 800–814 (2013).

8. Lo Presti, L. et al. Fungal effectors and plant susceptibility. Ann. Rev. Plant Biol. 66, 513–545 (2015).

9. Couto, D. & Zipfel, C. Regulation of pattern recognition receptor signalling in plants. Nat. Rev. Immunol. 16, 537–552 (2016).

10. Boller, T. & Felix, G. A renaissance of elicitors: perception of microbe-associated molecular patterns and danger signals by pattern-recognition receptors. Annu. Rev. Plant Biol, 60, 379–406 (2009).

11. Jones, J.D.G. & Dangl, J.L. The plant immune system. Nature, 444, 323–329 (2006).

12. Pegg, G.F. & Brady, B.L. Verticillium Wilts. Wallingford, Oxfordshire: CABI Publishing (2002).

13. Depotter, J.R. et al. Verticillium longisporum, the invisible threat to oilseed rape and other brassicaceous plant hosts. Mol. Plant Pathol. 17, 1004–1016 (2016).

14. Dunker, S., Keunecke, H., Steinbach, P. & von Tiedemann, A. Impact of Verticillium longisporum on yield and morphology of winter oilseed rape (Brassica napus) in relation to systemic spread in the plant. J. Phytopathol. 156, 698–707 (2008).

15. Tzelepis, G., Bejai, S., Sattar, M.N., Schwelm, A., Ilback, J., Fogelqvist, J. & Dixelius, C. Detection of Verticillium species in Swedish soils using real-time PCR. Arch. Microbiol. 199, 1383–1389 (2017).

16. Fahleson, J., Hu, Q. & Dixelius, C. Phylogenetic analysis of Verticillium species based on nuclear and mitochondrial sequences. Arc. Microbiol. 181, 435–442 (2004).

17. Inderbitzin, P., Bostock, R.M., Davis, R.M., UsamI, T., Platt, H.W. & Subbarao, K.V. Phylogenetics and taxonomy of the fungal vascular wilt pathogen Verticillium, with the descriptions of five new species. Plos One 6, e28341 (2011).

18. Inderbitzin, P., Davis, R.M., Bostock, R.M. & Subbarao, K.V. Identification and differentiation of Verticillium species and V. longisporum lineages by simplex and multiplex PCR assays. Plos One 8, e65990 (2013).

19. Chen, Y.A. & Scheller, R.H. SNARE-mediated membrane fusion. Nat. Rev. Mol. Cell Biol. 2, 98–106 (2001).

20. Lev, S., Ben Halevy, D., Peretti, D. & Dahan N. The VAP protein family: from cellular functions to motor neuron disease. Trends Cell. Biol. 18, 282–290 (2008).

21. Fogelqvist, J., Tzelepis, G., Bejai, S., Ilback, J., Schwelm, A. & Dixelius, C. Analysis of the hybrid genomes of two field isolates of the soil-borne fungal species Verticillium longisporum. BMC Genom. 19, 14 (2018).

22. Matoba, Y., Katsube, Y., & Sugiyama, M. The crystal structure of prokaryotic phospholipase A_2_. J. Biol. Chem. 277, 20059–20069 (2002).

23. Soragni, E., Bolchi, A., Balestrini, R., Gambaretto, C., PercudanI, R., Bonfante, P. & Ottonello, S. A nutrient-regulated, dual localization phospholipase A2 in the symbiotic fungus Tuber borchii. EMBO J. 20, 5079–5090 (2001).

24. Murakami, M. & Kudo, I. Secretory phospholipase A_2_. Biol. Pharm. Bull. 27, 1158–1164 (2004).

25. Tian, L., Yu, J., Wang, Y. & Tian, C. The C2H2 transcription factor VdMsn2 controls hyphal growth, microsclerotia formation, and virulence of Verticillium dahliae. Fungal Biol. 121, 1001–1010 (2017).

26. Zhang, T. et al. (2016). Cotton plants export microRNAs to inhibit virulence gene expression in a fungal pathogen. Nature plants 2, 16153 (2016).

27. Santhanam, P. & Thomma, B.P. Verticillium dahliae Sge1 differentially regulates expression of candidate effector genes. Mol. Plant Microbe Interact. 26, 249–256 (2013).

28. Stergiopoulos, I. & de Wit, P.J.G.M. Fungal effector proteins. Annu. Rev. Phytopathol. 47, 233–263 (2009).

29. Joosten, M.H.A.J., Vogelsang, R., Cozijnsen, T.J., VErberne, M.C. & de Wit, P.J.G.M. The biotrophic fungus Cladosporium fulvum circumvents Cf-4-mediated resistance by producing unstable AVR4 elicitors. Plant Cell 9, 367–379 (1997).

30. van den Burg, H.A., Harrison, S.J., Joosten, M.H.A.J., Vervoort, J. & de Wit, P.J.G.M. Cladosporium fulvum Avr4 protects fungal cell walls against hydrolysis by plant chitinases accumulating during infection. Mol Plant Microbe Interact. 19: 1420–1430 (2006).

31. Collmer, A., LIndeberg, M., Petnicki-Ocwieja, T., Schneider, D.J. & Alfano, J.R. Genomic mining type III secretion system effectors in Pseudomonas syringae yields new picks for all TTSS prospectors. Trends Microbiol, 10: 462–469 (2002).

32. Zurbriggen, M.D., Carrillo, N. & Hajirezaei, M.R. ROS signaling in the hypersensitive response: when, where and what for?. Plant signal. Behav. 5: 393–396 (2010).

33. Wendling, C., Spehner, D., Nominé, Y., Giordano, F., Mathelin, C., Drin, G., Tomasetto, C., & Alpy, F. Identification of MOSPD2, a novel scaffold for endoplasmic reticulum membrane contact sites. EMBO reports. 19: e45453.

34. Shi-Kunne, X., van Kooten, M., Depotter, J.R.L., Thomma, B.P.H.J. & Seidl, M. F. 2019. The genome of the fungal pathogen Verticillium dahliae reveals extensive bacterial to fungal gene transfer. Genome Biol. Evol. 11: 855–868. (2019).

35. Miozzi, L., Balestrini, R., Bolchi, A., Novero, M., Ottonello, S. & Bonfante, P. Phospholipase A(2) up-regulation during mycorrhiza formation in Tuber borchii. New Phytol. 167: 229–238 (2005).

36. Cavazzini, D., Meschi, F., Corsini, R., Bolchi, A., Rossi, G.L., Einsle, O. & Ottonello, S.. Autoproteolytic activation of a symbiosis-regulated truffle phospholipase A(2). J. Biol. Chem. 288: 1533–1547 (2013).

37. Beres, S.B, et al. Genome sequence of a serotype M3 strain of group A Streptococcus: phage-encoded toxins, the high-virulence phenotype, and clone emergence. Proc. Nat.l Acad. Sci. USA. 99:10078–10083 (2002).

38. Berthelot, P, Attree, I., Plésiat, P., Chabert, J., de Bentzmann, S., Pozzetto, B., Grattard, F; Groupe d’etudes des septicémies à Pseudomonas aeruginosa. Genotypic and phenotypic analysis of type III secretion system in a cohort of Pseudomonas aeruginosa bacteremia isolates: evidence for a possible association between O serotypes and exo genes. J. Infect. Dis. 188: 512–518 (2003).

39. Rowland, O., Ludwig, A.A., Merrick, C.J., Baillieul, F., Tracy, F.E., Durrant, W. E., Fritz-Laylin, L., Nekrasov, V., Sjölander, K., Yoshioka, H., & Jones, J.D. Functional analysis of Avr9/Cf-9 rapidly elicited genes identifies a protein kinase, ACIK1, that is essential for full Cf-9-dependent disease resistance in tomato. The Plant Cell 17: 295–310 (2005).

40. Pireyre, M., & Burow, M. Regulation of MYB and bHLH transcription factors: a glance at the protein level. Mol Plant. 8: 378–388 (2015).

41. van den Burg, H.A., Tsitsigiannis, D.I., Rowland, O., Lo, J., Rallapalli, G., Maclean, D., Takken, F.L., & Jones, J.D. The F-box protein ACRE189/ACIF1 regulates cell death and defense responses activated during pathogen recognition in tobacco and tomato. The Plant Cell. 20: 697–719 (2008).

42. van der Hoorn, R. Plant proteases: from phenotypes to molecular mechanisms. Annu. Rev. Plant Biol. 59: 191–223 (2008).

43. Figueiredo, A., Monteiro, F., & Sebastiana, M. Subtilisin-like proteases in plant-pathogen recognition and immune priming: a perspective. Front. Plant Sci. 5: 739 (2014).

44. Grewal, S., Morrison, E.E., Ponnambalam, S. & Walker, J.H. Nuclear localisation of cytosolic phospholipase A2-alpha in the EA.hy.926 human endothelial cell line is proliferation dependent and modulated by phosphorylation. J. Cell Sci. 115: 4533–4543 (2002).

45. Froidure, S., Canonne, J., Daniel, X., Jauneau, A., Brière, C., Roby, D., & Rivas, S. AtsPLA2-alpha nuclear relocalization by the Arabidopsis transcription factor AtMYB30 leads to repression of the plant defense response. Proc. Natl. Acad. Sci. USA. 107:15281–15286 (2010).

46. Peters-Golden, M., Song, K., Marshall, T. & Brock, T. 1996. Translocation of cytosolic phospholipase A_2_ to the nuclear envelope elicits topographically localized phospholipid hydrolysis. Biochem. J. 318: 797–803 (1996).

47. Lev, S., Ben Halevy, D., Peretti, D., & Dahan, N. The VAP protein family: from cellular functions to motor neuron disease. Trends Cell. Biol. 18: 282–290. (2008).

48. Petersen, N.H.T., Joensen, J., McKinney, L.V., Brodersen, P., Petersen, M., Hofius, D., Mundy, J. Identification of proteins interacting with Arabidopsis ACD11. Plant Physiol. 166:661–666 (2009).

49. Saravanan, R.S., Slabaugh, E., Singh, V.R., Lapidus, L.J., Haas, T., Brandizzi, F. The targeting of the oxysterol-binding protein ORP3a to the endoplasmic reticulum relies on the plant VAP33 homolog PVA12. Plant J. 58:817–830 (2009).

50. Kim, H., O’Connell, R., Maekawa-Yoshikawa, M., Uemura, T., Neumann, U., Schulze-Lefert, P. The powdery mildew resistance protein RPW8.2 is carried on VAMP721/722 vesicles to the extrahaustorial membrane of haustorial complexes. Plant J. 79: 835–847 (2014).

51. Santos, M.F., Rappa, G., Karbanová, J, Kurth, T., Corbeil, D., Lorico, A. VAMP-associated protein-A and oxysterol-binding protein-related protein 3 promote the entry of late endosomes into the nucleoplasmic reticulum. J. Biol. Chem. 293: 13834–13848 (2018).

52. Rafiei, V., Najafi, Y., Vélëz, H., & Tzelepis, G. Investigating the role of a putative endolysin-like candidate effector protein in Verticillium longisporum virulence, Biochem. Biophys. Res. Commun. 629: 6–11 (2022).

53. Rafiei, V., Ruffino, A., Persson-Hoden, K., Tornkvist, A., Mozuraitis, R., Dubey, M. & Tzelepis, G. A Verticillium longisporum pleiotropic drug transporter determines tolerance to the plant host beta-pinene monoterpene. Mol. Plant Pathol. 23: 291–303 (2022).

54. Livak, K.J. & Schmittgen, T.D. (2001). Analysis of relative gene expression data using real-time quantitative PCR and the 2(T)(-Delta Delta C) method. Methods. 25: 402-408 (2001).

55. Letunic, I., Doerks, T. & Bork, P. (2009). SMART 6: recent updates and new developments. Nucleic Acids Res, 37: D229–D232.

56. Sperschneider, J., Dodds, P.N., Singh, K.B., & Taylor, J.M. ApoplastP: prediction of effectors and plant proteins in the apoplast using machine learning. New phytol. 217: 1764–1778 (2018).

57. Sperschneider, J., Catanzariti, A.M., DeBoer, K., Petre, B., Gardiner, D. M., Singh, K. B., Dodds, P.N., & Taylor, J.M. LOCALIZER: subcellular localization prediction of both plant and effector proteins in the plant cell. Sci Rep. 7: 44598 (2017).

58. Grigoriev, IV et al. “MycoCosm portal: gearing up for 1000 fungal genomes.” Nucleic Acids Res. 42: D699–704 (2014).

59. Kumar, S., Stecher, G., Li, M., Knyaz, C. & Tamura, K. MEGA X: Molecular evolutionary genetics analysis across computing platforms. Molecul. Biol. Evol. 35: 1547–1549 (2018).

60. Waterhouse, A., Bertoni, M., Bienert, S., Studer, G., Tauriello, G., Gumienny, R., Heer, F.T., de Beer, T.A.P., Rempfer, C., Bordoli, L., Lepore, R., Schwede, T. SWISS-MODEL: homology modelling of protein structures and complexes. Nucleic Acids Res. 46: W296–W303 (2018).

61. Nygren, H., Seppänen-Laakso, T., Castillo, S., Hyötyläinen, T., & Orešič, M. (2011). Liquid chromatography-mass spectrometry (LC-MS)-based lipidomics for studies of body fluids and tissues. Methods Mol. Biol. 708: 247–257.

62. Utermark, J., & Karlovsky, P. Genetic transformation of filamentous fungi by Agrobacterium tumefaciens. Protocol Exchange. http://www.nature.com/protocolexchange/protocols/427, (2008).

63. Bushnell, B. BBTools: A suite of fast, multithreaded bioinformatics tools designed for analysis of DNA and RNA sequence dData. Jt. Genome Inst. Berkeley, CA, USA (2018).

64. Andrews, S. FastQC: a quality control tool for high throughput sequence data. Available at: https://www.bioinformatics.babraham.ac.uk/projects/fastqc/ (2010).

65. Dobin, A., et al. 2013. STAR: Ultrafast universal RNA-seq aligner. Bioinformatics. 29:15–21 (2013).

66. Langmead, B., & Salzberg, S. L. Fast gapped-read alignment with Bowtie 2. Nat. Methods. 9: 357–359 (2012).

67. Liao, Y., Smyth, G.K., & Shi, W. FeatureCounts: an efficient general purpose program for assigning sequence reads to genomic features. Bioinformatics. 30: 923–930 (2014).

68. Love, M.I., Anders, S., & Huber, W. Differential analysis of count data - the DESeq2 package. Genome Biol. 15:10–1186 (2014).

69. Pertea, G., & Pertea, M. GFF utilities: GffRead and GffCompare. F1000 Research. 9 (2020).

70. Urban, M., et al. PHI-base: A new interface and further additions for the multi-species pathogen-host interactions database. Nucleic Acids Res. 45:604–610 (2017).

