## Supplementary figure 1 for "The *Verticillium longisporum* phospholipase VlsPLA_2_ is a virulence factor targets host nuclei and modulates plant immunity"

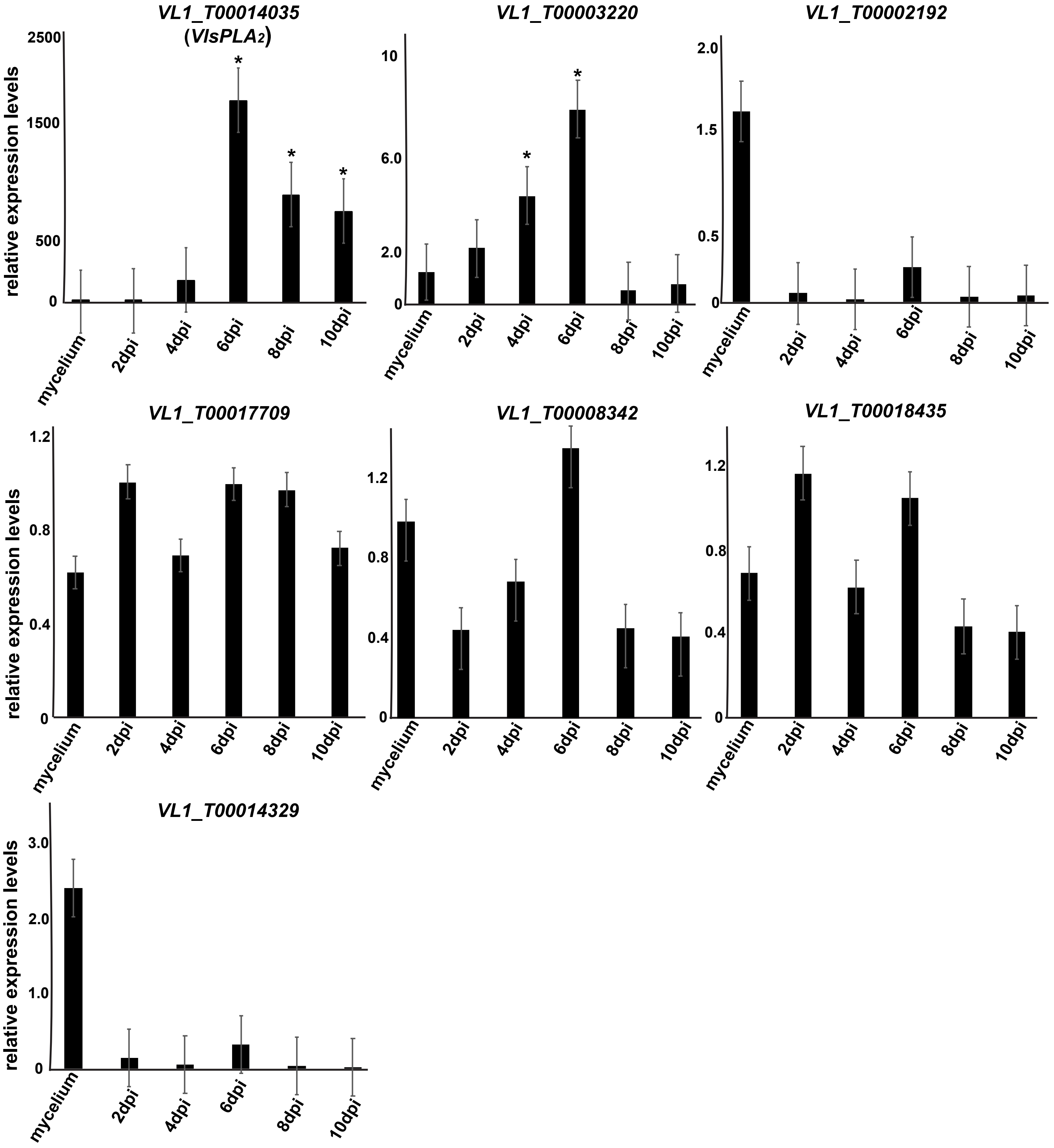


**Supplementary Fig. 1. Transcription profiles of *V. longisporum* genes encoding candidate effector proteins upon infection of *Brassica napus.*** Gene expression analysis was conducted according to $2^{-\Delta\Delta CT}$ method. Data were normalized using the expression levels of the reference gene, glyceraldehyde phosphate dehydrogenase (*GAPDH*). Error bars represent SE based on at least three biological replicates. Asterisks indicate statistically significant differences according to Students T' test (p value < 0.05).
