## Supplementary figure 2 for "The *Verticillium longisporum* phospholipase VlsPLA_2_ is a virulence factor targets host nuclei and modulates plant immunity"

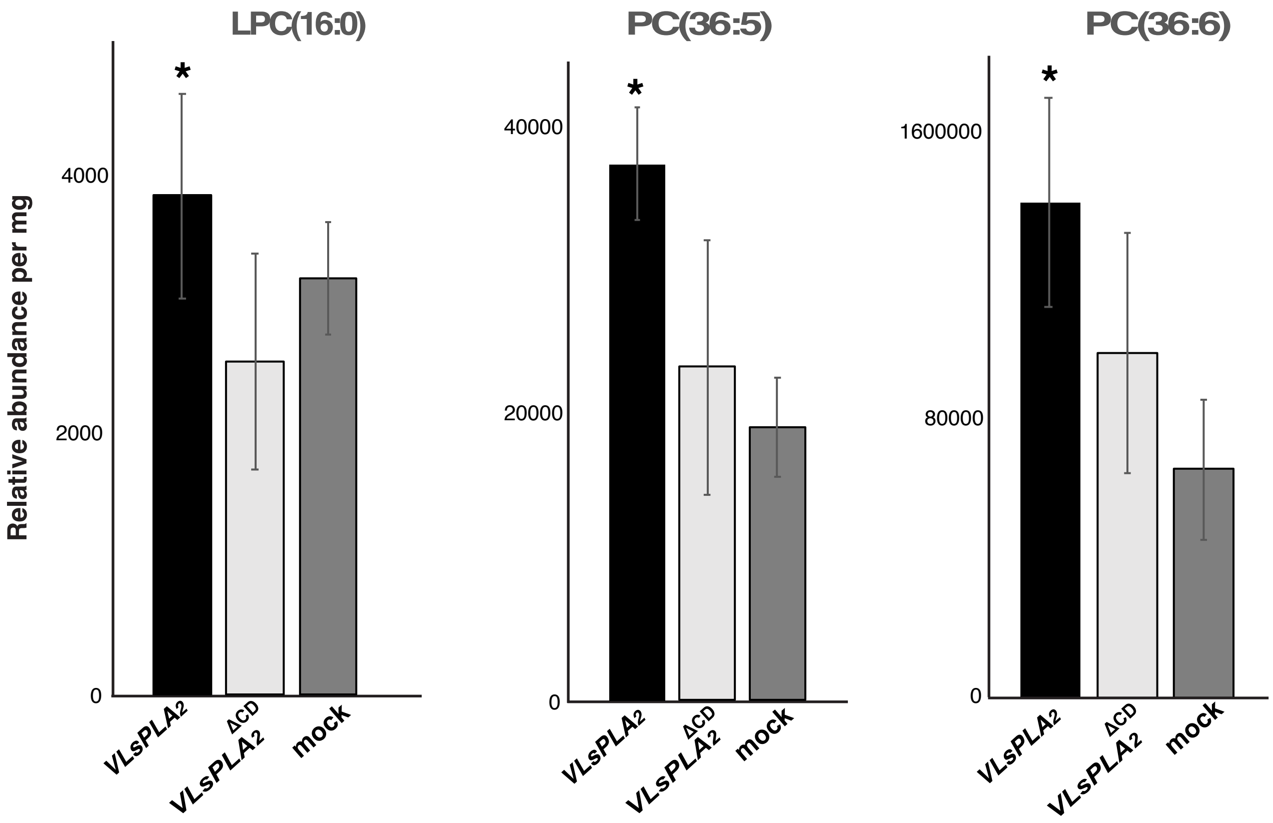


**Supplementary Fig. S2**. Phospholipid profiles on *N. benthamiana* transiently expressed either $VlsPLA_2{}^{WT}$ or $VlsPLA_2{}^{\Delta CD}$. Mocked-inoculated plants were used as control. Data depicted as a relative abundance of phospholipids per mg of dry plant tissue. Asterisks (*) indicate statistically significant differences according to the Student's T test ($p < 0.05$). Bars represent the standard error (SE) based on five biological replicates.
