## Supplementary figure 3 for "The *Verticillium longisporum* phospholipase VlsPLA_2_ is a virulence factor targets host nuclei and modulates plant immunity"

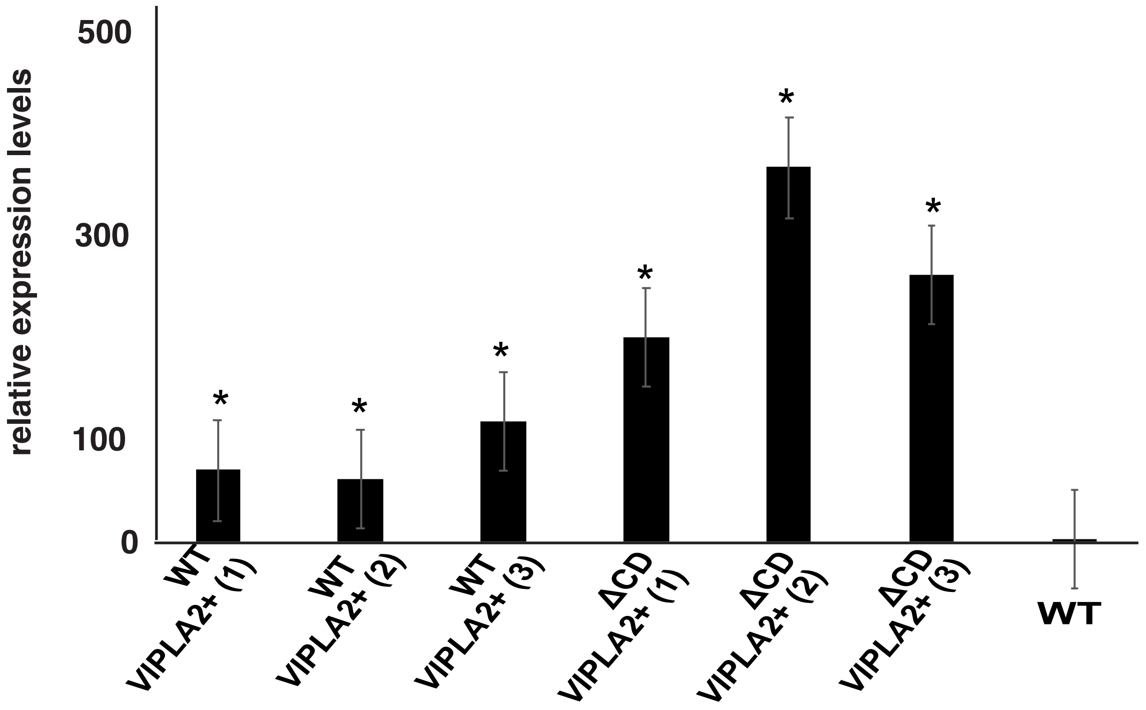


**Supplementary Fig. S3. Transcription analysis on *Verticillium longisporum* strains overexpressing either the functional active (VlsPLA$_2$+$^{WT}$) or the inactive (VlsPLA$_2$ + $^{\Delta CD}$) VlsPLA$_2$ phospholipase.** Gene expression analysis was conducted according to $2^{-\Delta\Delta CT}$ method. Data were normalized using the expression levels of the reference gene,glyceraldehyde phosphate dehydrogenase (*GAPDH*). Asterisks indicate statistically significant differences between the overexpression and WT strains according to the Students T' test (p value < 0.05).
