## Supplementary figure 4 for "The *Verticillium longisporum* phospholipase VlsPLA_2_ is a virulence factor targets host nuclei and modulates plant immunity"

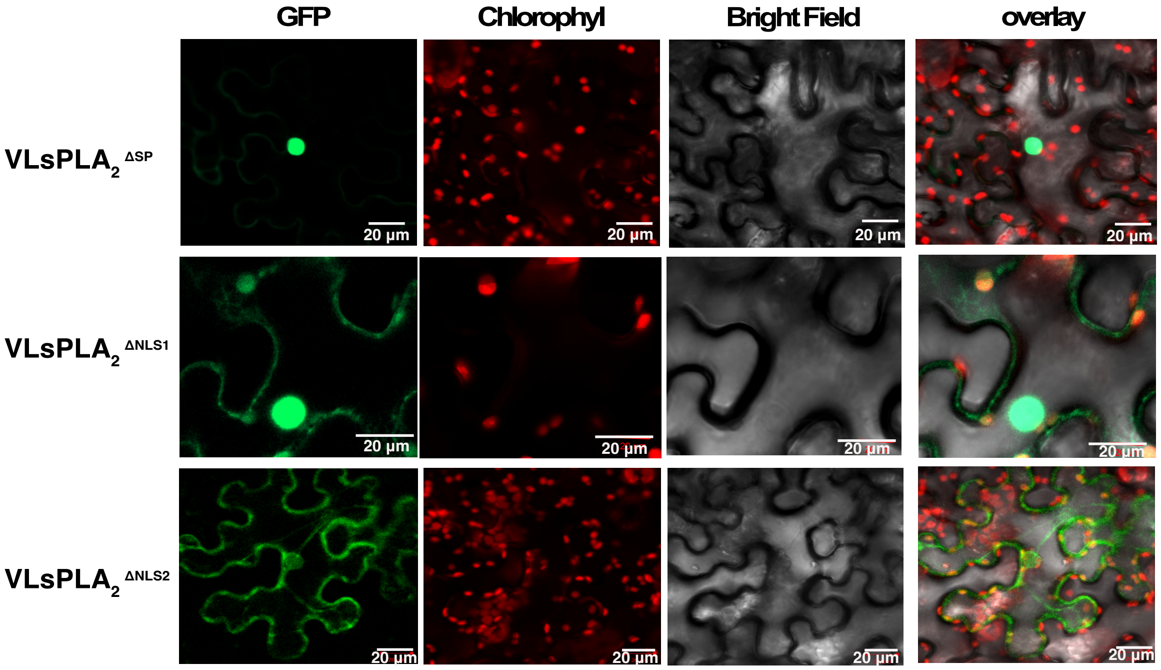


**Supplementary Figure S4.** Live-cell imaging of $VlsPLA_2$ mutants where signal peptide ($VLsPLA_2^{\Delta SP}$), or NLS1 ($VLsPLA_2^{\Delta NLS1}$), or NLS2 ($VLsPLA_2^{\Delta NLS2}$) were truncated. Mutants tagged with GFP at the C-terminal and Agro-infiltrated *N. benthamiana* leaves. The localization was monitored with a laser-scanning confocal microscope with a sequential scanning mode 48 hours post infiltration. The GFP and the chlorophyll were excited at 488 nm. GFP (green) and chlorophyll (red) fluorescent signals were collected at 505– 525 and 680–700 nm, respectively.
