## Supplementary figure 5 for "The *Verticillium longisporum* phospholipase VlsPLA_2_ is a virulence factor targets host nuclei and modulates plant immunity"

**a**

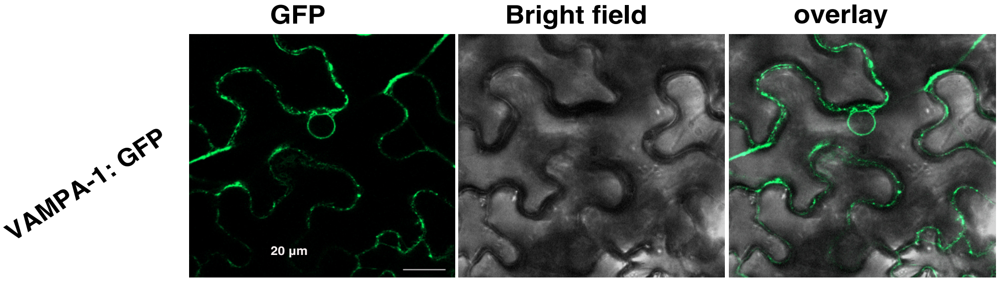


**b**

```
NbS00015616g0001.1      IQPWDSCFIRV-------------------------------------TLQAQKEYPLDM    73
NbS00038240g0008.1      IHPWDSCFIRVVHPNTDILLWFHLSRNMRHPFNGWFSYIVLNFVMSLVTLQAQKEYPPDM    103
NbS00060499g0005.1      IKPKSTYDFTV-------------------------------------TMQAQRTAPSDM    81
NbS00002269g0003.1      VLPRSRSNVTV-------------------------------------TMQPQTELPPDM    81
NbS00011956g0004.1      ------------------------------------------------TMQAQKEAPPDM    30
NbS00056068g0004.1      ------------------------------------------------TMQAQKEAPADM    32
NbS00026343g0007.1      VLPGSSCNVTV-------------------------------------TMQAQKEVPPDM    83
NbS00019865g0007.1      VLPGSSCNVTV-------------------------------------TMQAQKEAPPDM    83
NbS00032336g0008.1      VMPHSTCDVTV-------------------------------------TMQAQKEAPPDM    81
NbS00013560g0006.1      VLPRSTCDVIV-------------------------------------TMQAQKEAPADM    83
                                                                        *:* *   * **

NbS00015616g0001.1      QCKDKFLLQSTIVNND--IDELSPDTFNKESGRTVEESKLRVVYISPHSSPGHSED-FR-    129
NbS00038240g0008.1      QCKDKFLLQSTIVNND--SDELSPDTFNKDSGRTVEECKLRVVYISTHSSPGQFEDVFK-    160
NbS00060499g0005.1      QCKDKFLIQGTVVPFGTSEEEITPSMFNKDNRKYIEECKLRVVLVSPPNSPVLQPANGIS    141
NbS00002269g0003.1      QCKDKFLVQSVITNPGSTPKDINPDMFNKSGGNHVEECKLKVVYVSPPRKPSPVPEDSE-    140
NbS00011956g0004.1      QCKDKFLLQSVVASPGTTPKDITPEMFNKESGNHVDECKLRVAYVPPQ-PPSPVREGSE-    88
NbS00056068g0004.1      QCKDKFLLQSVVASSGTTAKDITPEMFNKEEGNHVEDCKLRVVYVPPQQPPSPVQEGSE-    91
NbS00026343g0007.1      QCKDKFLIQSVIAPSGTSNKDITTEMFNKEDGKVIDEFKLRVVYVP-ANPPSPVPEGSE-    141
NbS00019865g0007.1      QCKDKFLIQSVIAPSGTSNKDITTEMFNREDGKVIDEFKLRVVYVP-ANPPSPVPEGSE-    141
NbS00032336g0008.1      QCKDKFLLQSVVAGPGTTPKDITPEMFNKESGNHVDECKLRVAYVP-PQPPSPVREGSE-    139
NbS00013560g0006.1      QCKDKFLLQSVVATAGTSPKDITQEMFNKEPGRVVEECKLKVIYLPPPQPPSPVAEGSE-    142
                        *******:*..:.  .   .::. . **:.  . ::: **:*  :     *

NbS00015616g0001.1      -QSSDFTSN---------------------------------------------------    137
NbS00038240g0008.1      -QNSDVNSS---------------------------------------------------    168
NbS00060499g0005.1      KQGAPIETSMQQEKFPSGVENLPPAQTVGKNNKDIKFEEEIEVLDLGLSFAKNTESNTDE    201
NbS00002269g0003.1      -EWTSAGAS------------------E----------------------TDLQEHG---    156
NbS00011956g0004.1      -EGSSPRAS------------------I----------------------SENG------    101
NbS00056068g0004.1      -EGSSPRAS------------------V----------------------SENGTVNT--    108
NbS00026343g0007.1      -EGGSPRAS------------------L----------------------TEDESKSS--    158
NbS00019865g0007.1      -EGGSPRAS------------------L----------------------TEDESKSS--    158
NbS00032336g0008.1      -EGSSPRAS------------------I----------------------SENGAEFH--    156
NbS00013560g0006.1      -EGSSPGQS------------------L----------------------TENDSQNG--    159
                         .
```

**c**

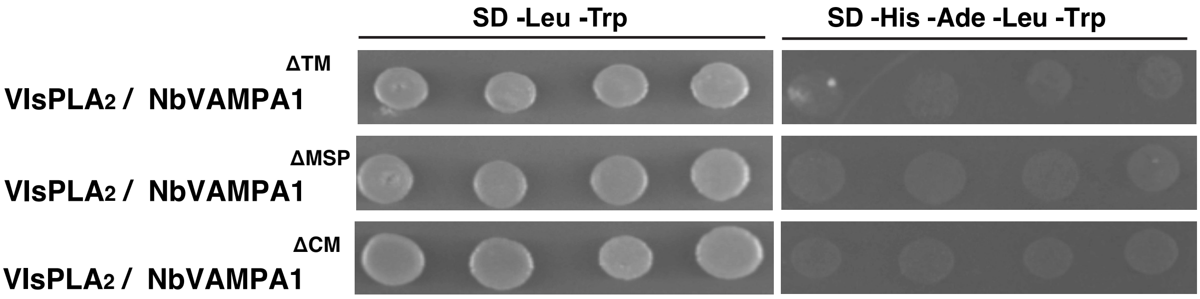


**Supplementary Fig. S5. Analysis of NbVAMPA1 protein. a.** Live-cell imaging of NbVAMPA-1 tagged with GFP at the N-terminal and Agro-infiltrated *N. benthamiana* leaves. The localization was monitored with a laser-scanning confocal microscope with a sequential scanning mode 48 hours post infiltration. The GFP was excited at 488 nm and collected at 505– 525 nm. **b.** Alignment of homologs to NbVAMPA1 in *N. benthamiana.* Identical sequences are marked with asterisks (*). **c.** Pairwise yeast-two-hybrid assays between $VlsPLA_2$ (used as a bait in pGBKT7 vector) and NbVAMPA1 mutated versions (used as a prey in pGADT7 vector). Growth of yeast cells on SD-4 (-His, -Ade, -Leu, -Trp) selective media represents protein–protein interaction and growth on SD-2 (-Leu, -Trp) media confirms yeast transformation
